# p53 pro-apoptotic activity is regulated by the G2/M promoting factor Cdk1 in response to DNA damage

**DOI:** 10.1101/2021.02.13.431094

**Authors:** Mireya Ruiz-Losada, Raul González, Ana Peropadre, Antonio Baonza, Carlos Estella

## Abstract

Exposure to genotoxic stress promotes cell-cycle arrest and DNA repair or apoptosis. These “life” or “death” cell fate decisions often rely on the activity of the tumor suppressor gene *p53*. Therefore, how *p53* activity is precisely regulated is essential to maintain tissue homeostasis and to prevent cancer development. Here we demonstrate that *Drosophila p53* pro-apoptotic activity is regulated by the G2/M kinase Cdk1. We find that cell cycle arrested or endocycle-induced cells are refractory to ionizing radiation induced apoptosis. We show that the p53 protein is not able to bind to and to activate the expression of the pro-apoptotic genes in experimentally arrested cells. Our results indicate that p53 genetically and physically interacts with Cdk1 and that p53 pro-apoptotic role is regulated by the cell cycle status of the cell. We propose a model in which cell cycle progression and p53 pro-apoptotic activity are molecularly connected to coordinate the appropriate response after DNA damage.

## Introduction

The ability of a cell to sense and to respond to DNA damage is essential to maintain its genetic material and tissue homeostasis. The DNA damage response (DDR) pathway has evolved in eukaryotes to preserve genomic integrity through a set of cellular responses that include cell cycle control, DNA repair, and apoptosis ^1^. Cell cycle regulation is an important response, as it allows the DNA repair mechanisms to prevent the incorrect transmission of genetic material, and therefore cancer susceptibility ^2^. Alternatively, if too much damage has been sustained, activation of cell death processes must occur to get rid of defective cells ^3^. Although the molecular mechanisms that control these cellular responses after DNA damage have been extensively studied separately, much less is known about how these processes are coordinated to maintain tissue homeostasis. Moreover, how the progression of the cell cycle impact in the ability of cells to activate the apoptotic response is mostly unexplored.

DNA lesions, such as double strand breaks (DSBs), are recognized by the MRE11– RAD50–NBS1 (MRN) protein complex that recruits and activates the ATM (ataxia-telangiectasia mutated) and ATR (ATM- and Rad3-Related) kinases ^4^. Activated ATM and ATR phosphorylate a number of substrates, such as the downstream kinases Chk1 and Chk2, which are responsible for the cell cycle checkpoint and apoptotic induction ^5^. Another ATM/ATR downstream protein target is the Histone H2AX (H2Av in *Drosophila*) that plays an essential role in the recruitment and accumulation of DNA repair proteins to sites of DSB damage ^6-8^.

Central in the DDR pathway is the tumor suppressor transcriptional protein p53, which can promote cell cycle arrest, DNA damage repair, apoptosis and senescence. Mutations in the p53 gene are strongly associated to cancer susceptibility and therefore it has been the focus of numerous studies ^9-11^. Mammalian p53 is activated by ATM and Chk2, and in turn p53 activates the expression of numerous target genes including the cell cycle regulator p21, the DNA repair protein Rad51 or the pro-apoptotic BCL-2 family members (*puma* and *noxa*) ^12-16^. The cellular context, timing and extent of the activation of the DDR pathway are responsible for the fate of the cell ^17^. In this sense, p53 activation and function requires a complex repertory of post-translational modifications and protein interactions ^18,19^. However, how p53 orchestrates these cell survival and cell death responses is largely unknown.

*Drosophila* has been widely used as a model to study the DDR as orthologs for many of the pathway components have conserved roles in regulating the responses after DNA damage ^20,21^. DSBs generated by ionizing radiation (IR) activate the ATM/ATR kinases, induce the phosphorylation of the H2Av, the cell cycle arrest through the ATR/Mei-41 and Chk1/Grapes axis and the apoptotic response mediated by the ATM/Tefu and Chk2/Mnk branch ^22-25^. As in mammals, *Drosophila* p53 is activated by Chk2 and triggers the activation of the pro-apoptotic genes *reaper, hid*, and *grim*. However, in *Drosophila* p53 is dispensable for IR-induced cell cycle checkpoint ^22,26-28^. While p53 is required for the rapid IR-induced apoptosis in imaginal discs, a p53-independent cell death that depends on c-Jun N-terminal kinase (JNK) pathway activation helps maintain genome integrity through the reduction of the number of aneuploid cells generated by IR 29-31.

In mammals and *Drosophila*, IR-induced apoptosis depends on the cell context and proliferation status of the cell ^32-39^. Most IR-resistant tissues are differentiated and non-proliferative cells. Identifying the molecular determinants that regulate apoptotic induction and its connection with the cell cycle machinery is essential to understand how cells coordinate the different responses after DNA damage ^40-42^.

Here, we use the *Drosophila* wing imaginal disc as a model to study how cell cycle progression impact in the ability of cells to induce DNA damage-induced apoptosis. We demonstrate that cell cycle arrested cells or endocycle-induced cells are insensitive to IR-induced apoptosis. We found that p53 activity and JNK pathway activation are compromised in cell cycle arrested cells. Specifically, we show that p53 is unable to bind to the regulatory regions of the pro-apoptotic genes in experimentally arrested cells. Consistent with this, we found that p53 and the G2/M promoting factor Cdk1 physically interact and that modification of Cdk1 activity influences p53 regulation of IR-induced apoptosis. We propose a model in which cell cycle progression and p53 activity are molecularly connected to coordinate the pro-apoptotic induction after DNA damage.

## Results

### Temporal dynamics of cell proliferation and apoptosis after DNA damage

To study the connection between cell cycle progression and DNA damage-induced apoptosis, we used the highly proliferative mono-layered of the *Drosophila* wing imaginal disc as a model. Exposure of wing imaginal cells to ionizing radiation (IR) generates DSB that induces a rapid cell cycle arrest and the activation of the apoptotic program ^22,25,27,43,44^. To monitor cell cycle dynamics, we used the Fly-FUCCI system (Fluorescence Ubiquitination-based Cell Cycle Indicator) and Fluorescence-activated cell sorting (FACS) to measure DNA content in combination with the mitotic marker phospho-Histone H3 (pH3). The FUCCI system is based on fluorochrome-tagged degrons from the Cyclin B (CycB, with Red Fluorescence Protein (RFP)) and E2F1 (with Green Fluorescence Protein (GFP)) proteins that are degraded during mitosis and G1 or at the onset of the S phase, respectively ^45^. Apoptotic cells were labeled using the initiator caspase reporter named DBS (for Drice-based sensor) ^46^. In the absence of apoptosis, the DBS sensor is restricted to the cellular membranes, however after inducing cell death it is translocated into the nucleus ^46^. As early as one hour after IR, there was a strong reduction in the number of mitotic cells as visualized by the absence of pH3, although no significant changes in the fraction of cells in G1 and G2 are observed (Fig. 1A-D). At this time point very few apoptotic cells were labelled with the DBS sensor. Three hours after IR, cells start to accumulate in G2 while a dramatic increase of apoptotic cells is detected (Fig. 1A-D). Six hours after IR, the G2/M mitotic arrest is lifted visualized by the recovery of pH3 positive cells and a high number of apoptotic cells are labeled (Fig. 1A-D). Interestingly, most of the apoptotic cells marked by the DBS sensor are not actively dividing, as they are pH3 negative.

**Figure 1:**
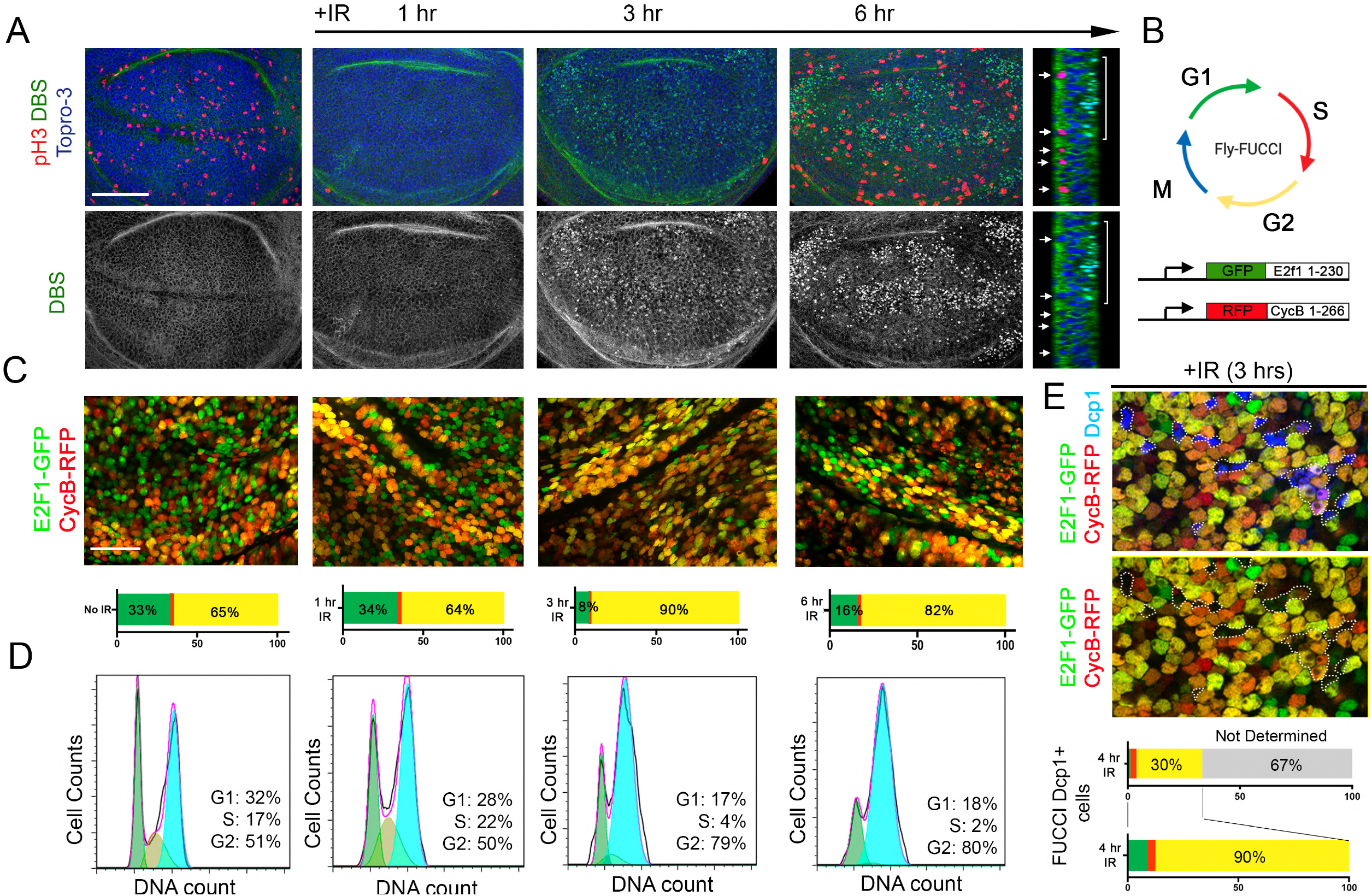
Temporal dynamics of cell cycle progression and apoptosis after IR. A) Third instar wing imaginal discs expressing the Dronc based sensor to mark apoptotic cells (DBS, in green) and stained with the mitotic marker pH3 (red) and Topro-3 (blue) to mark the nuclei. Representative examples of non-irradiated and irradiated discs dissected 1, 3 and 6 hrs after treatment. A Z-section of a wing imaginal disc dissected 6 hrs after IR is also show. Note that mitotic cells (red and arrows) are in a different plane and do not activate the DBS sensor (bracket). Scale bar: 50 um. B) Representation of the different phases of the cell cycle labeled with the Fly-FUCCI reporters (*UAS-GFP-E2F1 1-230* and *UAS-mRFP1-NLS-CycB 1-266*) and pH3 staining (blue). Cells are labeled in green (GFP+) in the G1 phase, in red during the S phase (RFP+), in yellow (GFP+RFP+) in the G2 phase and blue in mitosis. C) Third instar wing imaginal discs from the same treatment as in A, expressing the Fly-FUCCI transgenes under the *ap-Gal4* driver. The quantification of the number of cells that are GFP+, RFP+, and yellow (GFP+RFP+) for each condition is indicated below each image. Scale bar: 10 um. D) Bottom panels show cell cycle analysis by quantification of DNA content of control and irradiated wing imaginal discs dissected 1, 3 and 6 hrs after treatment. E) Wing imaginal disc cells dissected from a 3 hrs irradiated larvae expressing the Fly-FUCCI transgenes and stained with Dcp1 (blue). Dying cells are marked in blue and surrounded by a white dotted line. The quantification of the number of cells that are GFP+, RFP+ and yellow (GFP+RFP+) is indicated below. Apoptotic cells with very low or undetectable Fly-FUCCI reporters are considered as not determined. From those Dcp1+ cells with detectable Fly-FUCCI reporters, 90 % were in G2 phase.

We used the Fly-FUCCI system to visualize the phase of the cell cycle where IR-induced cells die. Apoptotic cells, identified by the effector caspase Dcp1, show very low or undetectable levels of the FUCCI reporters making it difficult to determine their exact cell cycle phase. However, in those cases in which we were able to detect the reporters, most of Dcp1 positive cells (90%) were in G2 (Fig. 1E).

These observations indicate that after IR, a G2/M arrest is rapidly activated followed by apoptosis induction.

### Cell cycle arrest block DNA damage induced apoptosis in wing imaginal discs

Next, we decided to explore in more detail the relationship between cell cycle progression and apoptotic induction after IR. To this end, we used the UAS/Gal4 system to express a battery of genetic tools in the *spalt* (*sal*) domain of the wing pouch to arrest cells at different phases of the cell cycle and study their effects on IR-induced apoptosis. We used the Fly-FUCCI system in combination with pH3 staining and FACS to precisely stage the phase of the cycle in which the cells are accumulated (Fig. 1B). Based on these markers, the expression of the p21 ortholog, *dacapo* (*dap*), the downregulation of CycE or E2F1 and the ectopic expression of an activated form of *Retinoblastoma* (*Rbf* ^*280*^) arrested cells in G1 (Fig. 2A-B and Fig. S1). A G2 and G2/M stalling was achieved by the knockdown of the Cdc25 phosphatase String (Stg) and the downregulation of the M-phase promoting factor Cdc2/Cdk1, respectively (Fig. 2A-B and Fig. S1). In addition, the expression of the Anaphase-Promoting Complex/Cyclosome (APC/C) binding protein Fizzy-related (Fzr/Cdh1) or the downregulation of CycA induces the transition from a mitotic cycle to an endocycle (Fig. 2A-B and Fig. S1). The endocycle is a modified cell cycle that alternates G and S phases without entering mitosis, resulting in large polyploid cells ^47^. All these cell cycle modifications are consistent with previous reports ^48-54^. Remarkably, IR-induced apoptosis in the wing disc, visualized by the effector caspase (Dcp1) staining, is strongly attenuated in cell cycle arrested cells and in endocycle-induced cells in the *sal* domain at 4 and 24 hrs after treatment (Fig. 2C and E and Fig. S2). This apoptosis attenuation in cell cycle arrested cells was confirmed by the TUNEL assay, which measures DNA fragmentation caused by cell death (Fig. S2A). Importantly, IR induced-apoptosis depends on the activity of the pro-apoptotic genes, as the expression of a UAS transgene that simultaneously inhibit the *rpr, hid*, and *grim* genes (UAS*-miRHG*) abolished cell death (Fig. 2D).

**Figure 2:**
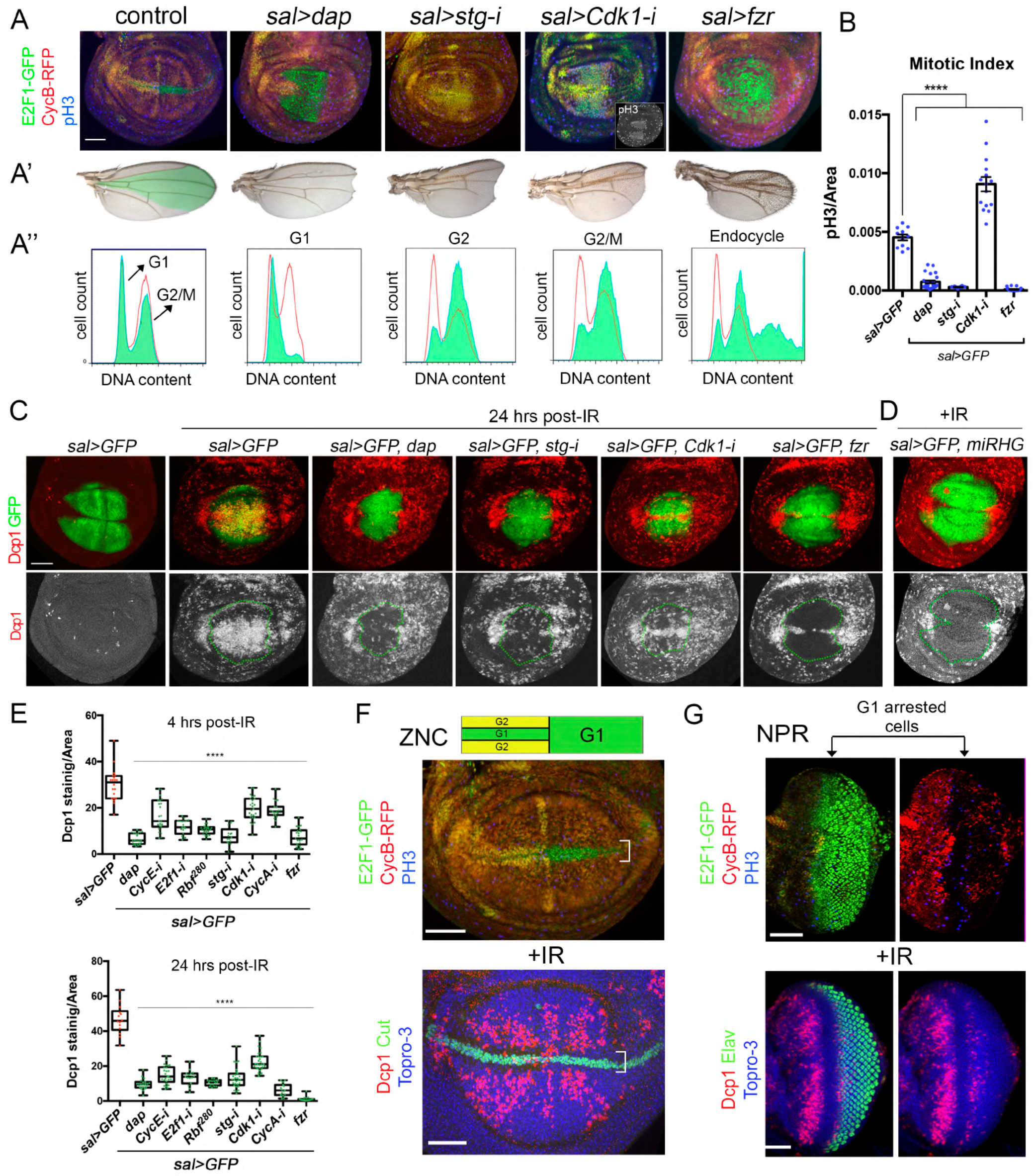
Cell cycle arrest and endocycle-induced cells attenuate IR-induced apoptosis. A) Cell cycle perturbations by the expression of *dap, stg-RNAi (stg-i), Cdk1-RNAi* (*Cdk1-i*) and *fzr* with the *sal-Gal4* (*sal>*) driver. The Fly-FUCCI system (*ubi-GFP-E2F11-230* and *ubi-mRFP1-NLS-CycB1-266*) and pH3 staining (blue) was used to visualize the cell cycle. The inset of the *sal>Cdk1-i* panel also shows the separate channel for the pH3 staining (white). The adult wing phenotypes of these experimentsare shown in A’ where the *sal* domain is colored in green in the control. Cell cycle profiles of dissociated wing imaginal discs expressing the indicated cell-cycle regulators and GFP in the *sal* domain is shown in A’’. Red profiles correspond to control GFP negative cells and green profiles belong to GFP positive cells in control and cell cycle perturbed cells. B) Mitotic index measured as the number of pH3 positive cells per area in the *sal* domain of control (*sal>GFP*) and in cell cycle arrested cells and endocycle-induced cells. n>11 discs per genotype. Error bars indicate standard error of the man (SEM). **** P value <0,0001 by one-way ANOVA when compared the mean of each column with the mean of the control. C-D) GFP (green) and Dcp1 staining (red and white) in wing imaginal discs expressing the indicated transgenes by the *sal-Gal4* driver in control discs and irradiated discs analyzed 24 hrs later. Below each panel, the Dcp1 channel is shown and the *sal* domain is outlined by green dotted lines. E) Quantification of Dcp1 staining in the *sal* domain in wing imaginal discs expressing the indicated transgenes by the *sal-Gal4* in irradiated discs analyzed 4 and 24 hrs after treatment. Error bars indicate the minimum and maximum point for each genotype. Individual wing discs measurements are shown. n>15 discs per genotype. **** P value <0,0001 by one-way ANOVA when compared the mean of each column with the mean of the control (*sal>GFP*). Scale bar: 50 um. F) Scheme of the zone of non-proliferating cells (ZNC) of the wing imaginal disc and a wing carrying the Fly-FUCCI transgenes (*ubi-GFP-E2F11-230* and *ubi-mRFP1-NLS-CycB1-266*) and stained with pH3 (blue) to label cells in G1 (green), G2 (yellow), S (red) and M (blue) phases. Below, a third instar wing imaginal disc from irradiated larvae dissected 4 hrs later. Dcp1 is in red, Cut in green and Topro-3 in blue. The brackets indicate the ZNC. Non-proliferating region (NPR) of the eye-antenna imaginal disc stained with Fly-FUCCI and pH3. Cells in G1 (green), S (red), G2 (yellow) and M (blue) phases are labeled. Below, an eye-antenna imaginal disc from irradiated larvae dissected 4 hrs later. Dcp1 is in red, Elav in green and Topro-3 in blue. Note the absence of Dcp1 staining in the NPR (arrows). See also Figures S1 and S2.

Next, we analyzed the apoptotic response to IR in cells that are developmentally arrested in different phases of the cell cycle. We focus on the zone of non-proliferating cells (ZNC) that coincides with the wing margin of the disc and in a specific region of the eye disc, known as the non-proliferative region (NPR), where cells are arrested in G1 ^55,56^(Fig. 2F and G). Our results confirm previous reports that showed that IR-induced apoptosis is strongly suppressed in any of these developmentally arrested regions (Fig. 2F and G) ^32,57^.

These results demonstrated that the apoptotic response after IR is compromised in developmentally and experimentally induced cell cycle arrested cells.

### Analysis of the DNA damage response pathway in cell cycle arrested and endocycle-induced cells after IR

To analyze whether the attenuation of IR-induced apoptosis in cell cycle arrested cells is caused because the activity of DDR pathway is compromised, we study ATM/ATR activity through pH2AV staining. We used the *hh*-*Gal4* driver in combination with the *tub-Gal80*^*ts*^ (*hh*^*Gal80*^*>*) to spatially and temporally arrest cells by the expression of *dap, stg-i* or *Cdk1-i* in the posterior compartment and stained for pH2Av. In addition, we also induced the endocycle by *fzr* forced expression in posterior cells. In a control disc 4 hrs after IR, an elevated number of strong nuclear pH2Av foci are readily detected accompanied with an increase in the overall staining of wing cells (Fig. 3A). These strong pH2Av foci are always associated with apoptotic cells in control and irradiated discs (Fig. S3A) ^58^. However, not all apoptotic cells are pH2AV positive. Consequently, the number of pH2Av foci in irradiated discs was reduced when the apoptotic pathway is compromised by the expression of the UAS*-miRHG* or by the knockdown of p53 levels (Fig. 3A, B and data not shown). In irradiated cell cycle arrested or endocycle-induced cells the number of pH2Av foci was also dramatically decreased when compared to the anterior compartment of the same disc or an irradiated control disc (Fig. 3A and B). Notably, the background levels of pH2Av staining were also slightly reduced in irradiated cell cycle arrested cells, especially for the *dap* or *stg-i* expressing discs, when compared to the corresponding anterior control cells (Fig. 3A).

**Figure 3:**
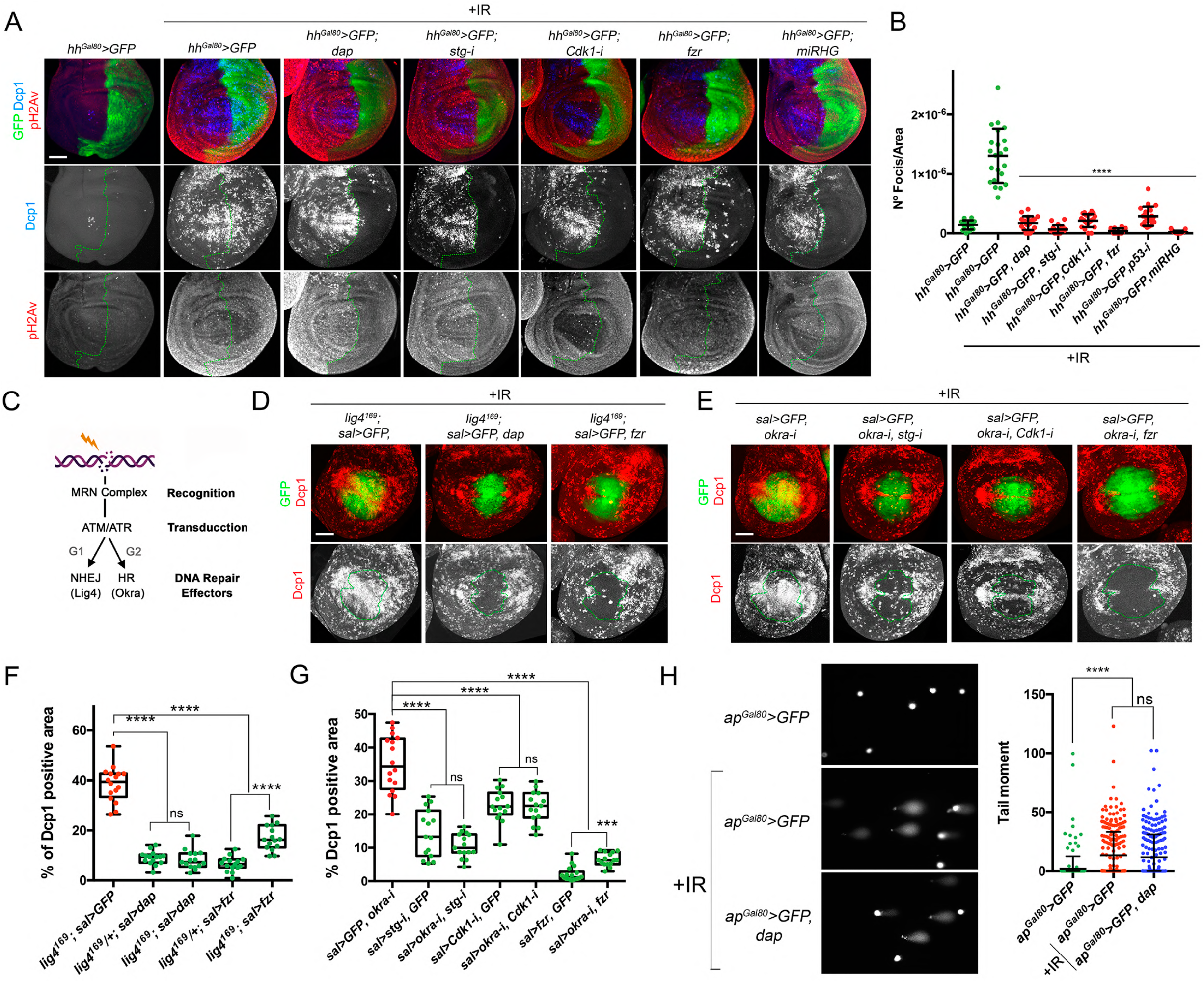
DDR pathway analysis in cell cycle arrested and endocycle-induced cells of irradiated wing imaginal discs. A) Wing imaginal discs expressing the indicated transgenes by the *hh-Gal4, tub-Gal80*^*ts*^ (*hh*^*Gal80*^*>*) subjected to IR and analyzed 4 hrs after treatment. Imaginal discs are stained with pH2Av (red and white), Dcp1 (blue and white) and GFP (green). The antero-posterior compartment boundary is marked by a green dotted line. Wing discs were dissected from larvae raised at 29°C for 24 hrs. Scale bar: 50 um. B) Scatter plots showing the number of pH2Av foci per area from control (non-irradiated) and irradiated imaginal discs expressing the different transgenes as in A. The average, standard deviation (SD) and individual measurements are shown. n>14 discs per genotype. **** P value <0,0001 by one-way ANOVA when compared the mean of each column with the mean of the control (*sal>GFP* IR). C) Simplified representation of the DDR pathway and the DNA repair mechanisms activated by DSB. D) Wing imaginal discs analyzed 24 hrs after IR treatment expressing the indicated transgenes by the *sal-Gal4* driver in a *lig4*^*169*^ mutant background. *dap* and *fzr* expression were selected as they arrested cells in G1 or induced the endocycle, respectively. GFP is green and Dcp1 staining is red or white. Below each panel, the Dcp1 channel is shown and the *sal* domain is outlined by green dotted lines. Scale bar: 50 um. E) Wing imaginal discs analyzed 24 hrs after IR treatment expressing the indicated transgenes by the *sal-Gal4. stg-i* and *Cdk1-i* arrested cells in G2 and G2/M, respectively while *fzr* induced the endocycle. GFP is green and Dcp1 staining is red or white. Below each panel, the Dcp1 channel is shown and the *sal* domain is outlined by green dotted lines. Scale bar: 50 um. F-G) Quantification of Dcp1 staining in the *sal* domain of wing imaginal discs of the corresponding genotypes shown in D and E. Error bars indicate the minimum and maximum point for each genotype. Individual wing discs measurements are shown. n>15 discs per genotype. **** P value <0,0001, *** P value <0,001 by one-way ANOVA when compared the mean of each experiment with the mean of the corresponding control. ns, not significant. H) Representative images of individual wing disc cells expressing *GFP* or *GFP* and *dap* under the *ap-Gal4, Gal80*^*ts*^ driver (*ap*^*Gal80*^*>*) for 24 hrs in control and irradiated discs and subjected to the Comet assay 5 hrs later. The tail DNA moment quantification for wing cells of the indicated genotypes is shown. Each dot represents a single cell. >250 comets were analyzed for each genotype and condition. Statistically significant differences based on Student’s t test are indicated: ****P < 0,0001 and not significant (ns). See also Figure S3.

The reduction of pH2Av staining after IR could suggest that DNA lesions are being repaired in cells that have been permanently arrested or shifted towards an endocycle, and therefore explains the attenuation of the apoptotic pathway. To test this possibility, we knocked down specific components of the DNA repair machinery in cell cycle arrested or endocycle-induced cells (Figure 3C). DNA lesions are recognized by the Mre11-Rad50-Nbs1 (MRN) complex and the repair of DSB is mediated by ATM/ATR through two different mechanisms: an error-prone non-homologous end-joining (NHEJ) and an error-free homologous recombination (HR). NHEJ is executed by Lig4 and preferentially operates in G1 while HR takes place in S and G2 where the sister chromatid is used as a template by DNA repair proteins such as Rad54/Okra ^21,59,60^. Importantly, the ability to attenuate apoptosis after IR in cell cycle arrested and endocycle-induced cells is maintained even when the recognition of DSB or the NHEJ and HR mechanisms are compromised (Figure 3D-G and Fig. S3). Similar results were obtained in a *mei41/ATR* mutant background or in knockdowns of Tefu/ATM (Fig. S3).

In addition, we measured IR-induced DNA lesions using the comet assay in control and cell cycle arrested cells. The comet assay consists in a single-cell gel electrophoresis test to visualize DNA damage. Comet’s DNA-tail provides information about the extent of DNA lesions and is represented as the tail moment. After IR, DNA damage (tail moment) is induced in control proliferating imaginal cells (*ap-Gal4, tub-Gal80*^*ts*^*>GFP*). However, no differences were observed in total DNA damage induced by IR in control discs *vs* G1 arrested discs (*ap-Gal4, tub-Gal80*^*ts*^ *>GFP, dap*) analyzed 5 hrs after treatment (Fig. 3H). From these results we conclude that IR produces comparable DNA lesions in proliferating and cell cycle arrested cells.

In summary, all together these results strongly suggest that the inhibition of IR induced-apoptosis observed in cell cycle arrested and endocycle-induced cells is not due to the prolonged repair of DNA lesions.

### Cell cycle arrested and endocycle-induced cells block apoptosis upstream of the pro-apoptotic genes

In order to understand a what level of the DDR pathway the cell cycle arrested and endocycle-induced cells block IR-induced apoptosis, we first follow the activation of the pro-apoptotic gene *hid* in irradiated wing imaginal discs. Four hours after IR, the levels of a GFP-tagged form of the Hid protein are clearly increased compared to non-irradiated disc (Fig. 4A). However, Hid-GFP levels are strongly downregulated in cell cycle arrested or endocycle-induced cells of irradiated wing imaginal discs (Fig. 4A). We confirmed the transcriptional repression of the *hid* gene in these cell cycle altered cells using a specific *hid cis*-regulatory module that is strongly induced after IR ^61,62^ (Fig. S4A). These results suggest that the inhibition of apoptosis is upstream of the pro-apoptotic genes. To confirm this hypothesis, we ectopically expressed the pro-apoptotic genes *hid* or *rpr* in the *sal* domain of the wing pouch in control and cell cycle arrested cells. In control discs, the expression of any of these pro-apoptotic genes strongly induced the activation of Dcp1. Accordingly, the cell cycle arrest or the induction of the endocycle overall did not have a strong impact on the ability of *rpr* or *hid* to induce apoptosis, confirming our previous results that demonstrated that the blockage of apoptosis must be upstream of the pro-apoptotic genes (Fig. 4B and Fig. S4B). *rpr* and *hid* are both regulated by p53 and the JNK pathway in response to genotoxic stress signals such as IR ^27,62-64^. Given the reported role of JNK in the induction of apoptosis after IR we monitored the activity of the pathway with a transcriptional reporter named *JNK*^*REP*^, which is induced in response to JNK signalling ^63,65^. In control discs, the *JNK*^*REP*^ is strongly activated after IR, however, in cell cycle arrested cells or endocycle-induced cells, the levels of the *JNK*^*REP*^ are remarkably downregulated (Fig. 4C). Consequently, the expression of a constitutive active form of JNKK Hemipterous (*hep*^*CA*^) induced the apoptotic pathway even in cells where the cell cycle has been arrested or shifted towards an endocycle (Fig. 4D).

**Figure 4:**
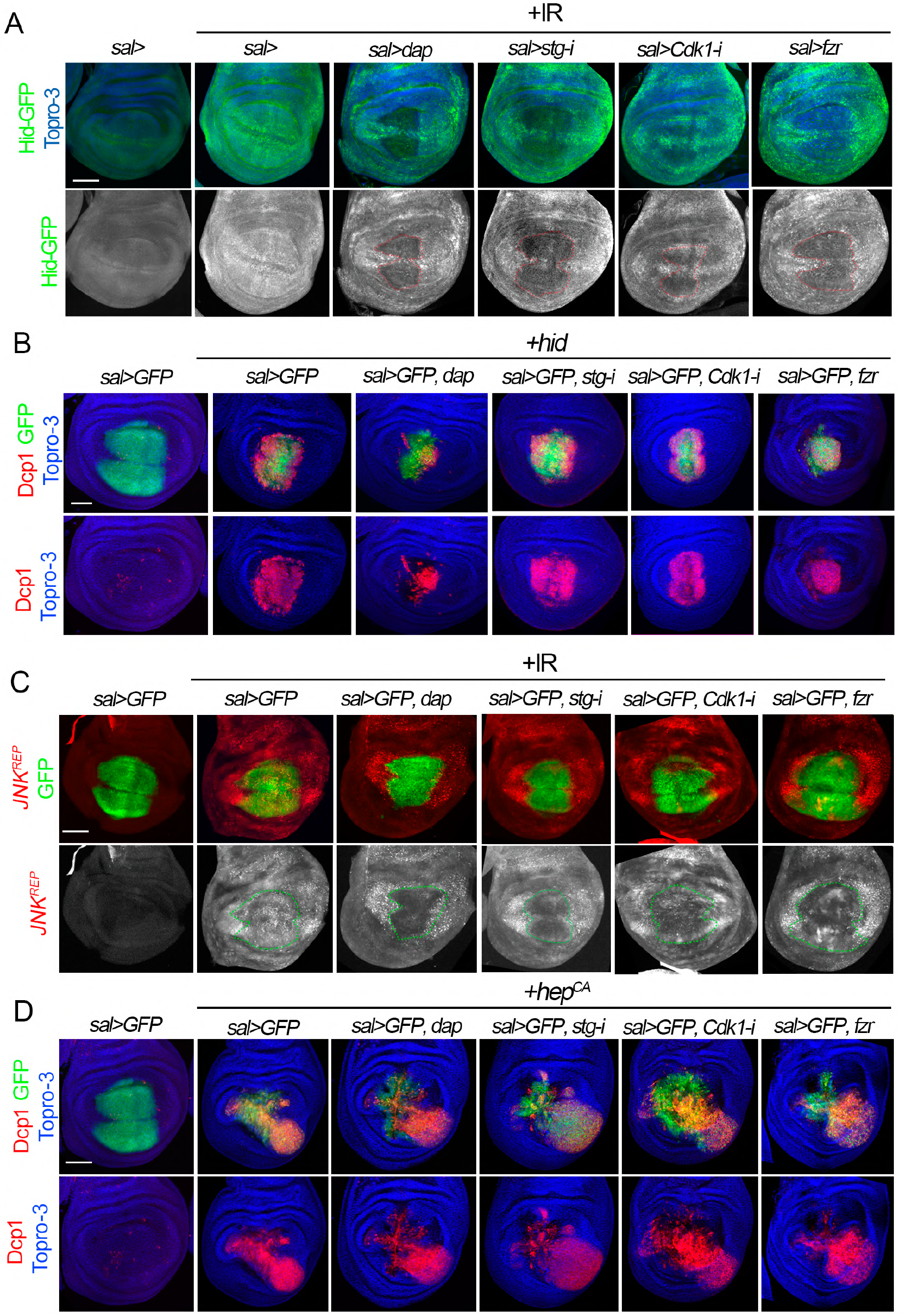
Analysis of Hid, Dcp1 and JNK pathway activation in cell cycle arrested and endocycle-induced cells. A) Third instar wing imaginal discs of the indicated genotypes that also express a GFP-tagged form of Hid (green and white). Note that after 4 hrs of IR, the levels of Hid increase significantly compared to a non-irradiated control. In cell cycle arrested and endocycle-induced cells in the *sal* domain, Hid levels are downregulated. Hid separate channel is shown below (white) and the *sal* domain is marked by red dotted lines. Topro-3 (blue) marks the nuclei. B) Wing imaginal discs that express in the *sal* domain (green) the pro-apoptotic gene *hid* and the corresponding cell cycle alterations indicated in the genotypes above each panel. Dcp1 in red, GFP in green and Topro-3 in blue. Note that although the ability of Hid to induce cell death is overall not altered in cell cycle arrested cells, we observed a reduction in Dcp1 activation in cells expressing *dap* when compared to the control. C) Wing imaginal discs expressing the indicated transgenes under the *sal-Gal4* driver and the JNK pathway activity reporter (*JNK*^*REP*^, red). Note that the *JNK*^*REP*^ is not active in the wing pouch, however it is strongly induced 24 hrs after IR. *JNK*^*REP*^ separate channel (white) is shown below each panel and the *sal* domain is marked by green dotted lines. D) The activation of the JNK pathway through the expression of *hep*^*CA*^ in the *sal* domain induced a strong activation of Dcp1 (red) even in cells where the cell cycle is blocked or shifted to an endocycle. The different combination of transgenes expressed with the *sal-Gal4* driver is indicated. Dcp1 in red, GFP in green and Topro-3 in blue. In A, B, C and D the scale bar is 50 um. See also Figure S4.

In summary, these results suggest that the blockade of IR induced apoptosis observed in cell cycle arrested cells is upstream of the pro-apoptotic genes.

### The pro-apoptotic activity of p53 is compromised in cell cycle arrested cells and endocycle-induced cells

Initial IR-induced cell death requires p53 activity, which activates the JNK pathway and the expression of the apoptotic genes ^27,62-64^. Our observations suggest that non-proliferating cells and endocycle-induced cells attenuate IR-induced apoptosis upstream of the pro-apoptotic genes. Therefore, we decided to study the connection between cell cycle progression and p53 regulation of IR-induced apoptosis. *Drosophila* has a single *p53* family member that encodes for four isoforms (A, B, C, and E). p53-A, also known as ΔNp53 because it lacks 110 amino acid in its N-terminal TAD when compared to the longest p53-B isoform, is the most abundant isoform in imaginal discs and the one responsible for the IR-induced apoptosis ^41,66,67^. Therefore, we analyzed p53 protein levels in cells of the wing imaginal disc that have been arrested at different phases of the cell cycle or that have been shifted to an endocycle. p53 is readily detected in wing imaginal discs by a specific antibody (Fig. 5A). No significative changes in p53 levels or protein localization were observed in experimentally arrested cells both in non-irradiated and irradiated wing discs (Fig. 5A and Fig. S5). Next, we tested whether the block on cell proliferation could have an impact on the ability of p53 to activate the expression of *hid* and to induce apoptosis.

**Figure 5:**
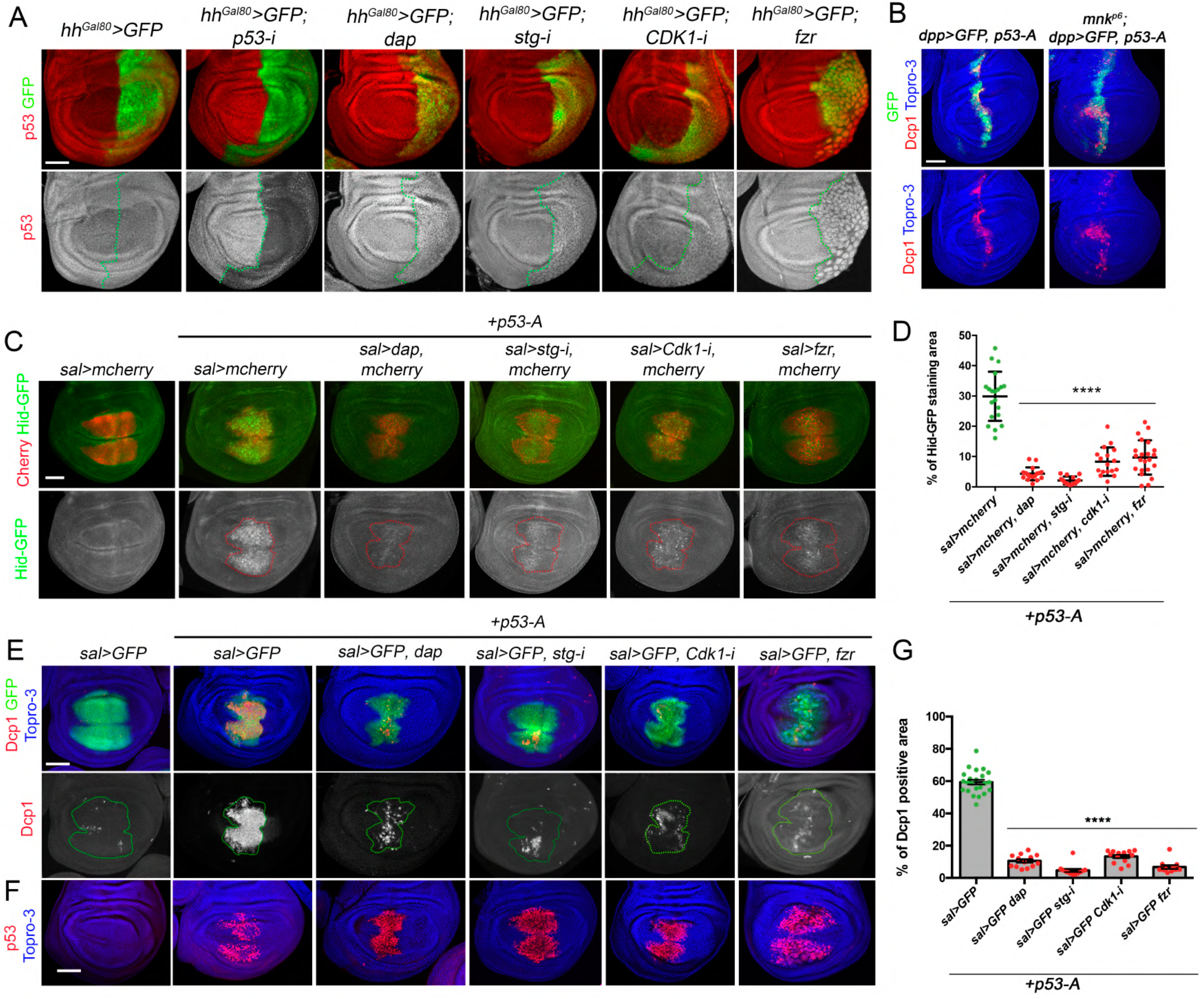
Analysis of p53 protein levels and activity in cell cycle arrested and endocycle-induced cells. A) Third instar wing imaginal discs expressing the indicated transgene in the posterior compartment by the *hh-Gal4, tub-Gal80*^*ts*^ (*hh*^*Gal80*^*>*) driver and stained for p53 (red and white) and GFP (green). Separate channel for p53 is shown below each image. Note that the expression of the *p53-i* eliminates p53 staining. The antero-posterior compartment boundary is marked by a green dotted line. Wing discs were dissected from larvae raised at 29°C for 24 hrs. B) Wing imaginal discs expressing *p53-A* under the *dpp-Gal4, UAS-GFP* (*dpp>GFP*) driver in a wild type and *mnk*^*p6*^ mutant background stained for Dcp1 (red), GFP (green) and Topro-3 (blue). C) Wing imaginal discs expressing the GFP-tagged form of Hid (green and white) and the indicated transgenes under the *sal-Gal4, UAS-mcherry* (red) driver. Separate channel for Hid-GFP is shown below each image. A dotted red line marks the *sal* domain. All the images were taken keeping the same confocal settings. D) Scatter plots showing the quantification of Hid-GFP staining in the *sal* domain from disc expressing *p53-A* in control (*sal>mcherry*) and cell cycle arrested cells as indicated in C. The average, standard deviation (SD) and individual measurements are shown. n>15 discs per genotype. **** P value <0,0001 by one-way ANOVA when compared the mean of each column with the mean of the control (*sal>mcherry, p53-A*). E-F) Wing imaginal discs expressing the corresponding transgenes under the *sal>GFP* driver. Imaginal discs were stained for Dcp1 (red and white), GFP (green), Topro-3 (blue) in E and for p53 (red) and Topro-3 (blue) in F. Separate channel for Dcp1 is shown below each image. A dotted red line marks the *sal* domain. G) Quantification of Dcp1 staining in the *sal* domain of wing imaginal discs from the genotypes presented in E. Error bars indicate SEM. n>11 disc per genotype. **** P value <0,0001 by one-way ANOVA when compared the mean of each column with the mean of the control (*sal>GFP, p53-A*). In A, B, C, E and F the scale bar is 50 um. See also Figure S5.

Overexpression of *p53-A* in the wing imaginal disc strongly activated *hid* expression and induced apoptosis (Figure 5C and E). Importantly, the expression of *p53-A* in cells that are simultaneously arrested or shifted towards an endocycle have a dramatically reduced ability to activate the apoptotic program as visualized by *hid* activation and Dcp1 staining (Fig. 5C-G). These effects were not due to the degradation of p53, as the levels the protein are comparable in *p53-A* overexpression in control cycling cells and arrested cells (Fig. 5F).

One possibility that could explain the decrease apoptotic activity of p53-A in cell cycle arrested and endocycling-induced cells could be that the DDR pathway is not properly activated, as we have reported by the analysis of pH2Av foci in these cells. Following IR, the ATM/Tefu checkpoint kinase phosphorylates several substrates such as H2Av and Chk2/Mnk that in turn phosphorylates p53. Like *p53* mutants, *mnk* mutants, the *Drosophila* ortholog of *Chk2*, are defective in IR-induced apoptosis ^25,26,68^. Importantly, the overexpression of *p53-A* in an *mnk/Chk2* mutant background induced apoptosis to the same extent as in a wild type background (Fig. 5B). These results indicate that the reduced ability of p53-A to trigger apoptosis in cell cycle arrested cells is not due to defects on upstream activation by Chk2.

### Cell cycle arrested cells block p53-A ability to bind to the Irradiation Responsive Enhancer Region (IRER) of the pro-apoptotic genes

It has been proposed that endocycling cells of the salivary glands are refractory to p53 induction of apoptosis by two possible mechanisms ^41^: First, a proteasome dependent p53 protein degradation and second, an epigenetic silencing of the pro-apoptotic genes at the H99 locus that blocks the activity of p53 to induce cell death. We thought that cell cycle arrested and endocycle-induced cells in wing imaginal discs that are refractory to p53-A apoptotic induction could employ a similar mechanism. Previously, we have shown that p53 protein levels are comparable in cycling *vs* non-cycling wing imaginal cells. Therefore, we decided to test whether non-proliferating cells could induce an epigenetic blockage at the level of the pro-apoptotic genes that prevents their activation by p53. An H99 intergenic regulatory region has been identified that mediates p53 induction of *rpr* and *hid* after IR in embryos and imaginal discs ^42,69^ (Fig. 6A). This region, named <u>I</u>rradiation <u>R</u>esponsive <u>E</u>nhancer <u>R</u>egion (IRER) contains a p53 responding element (p53^RE^) and its chromatin accessibility is epigenetically regulated 42,69. In endocycling salivary gland cells and late stage embryos, repressive chromatin marks have been found at the loci of the pro-apoptotic genes that correlate with lower p53 binding and IR-induced apoptosis ^41,42^. We take advantage of a chromatin state sensor, named IRER{ubi-DsRed}, that reflects the DNA accessibility at the IRER locus 69 (Fig. 6A). We reasoned that if cell cycle arrested and endocycle-induced cells prompted a repressive chromatin change at the IRER region that blocks p53 activation of the pro-apoptotic genes, we should observed a decrease in the IRER{ubi-DsRed} activity. First, we confirmed the responsiveness of the IRER{ubi-DsRed} sensor to p53-A overexpression or after the knockdown of the repressive histone methyltransferase Su(var)3-9 (Fig. 6B). Second, we tested IRER{ubi-DsRed} activity in cell cycle blocked cells and endocycle-induced wing cells. No changes in the IRER sensor activity were observed in these cells, suggesting that chromatin accessibility at this region is not affected by the cell cycle status of the cell (Fig. S6). In contrast, we found that p53-A ability to open the IRER is strongly reduced when p53-A is overexpressed in arrested or endocycle-induced wing cells (Fig. 6C).

**Figure 6:**
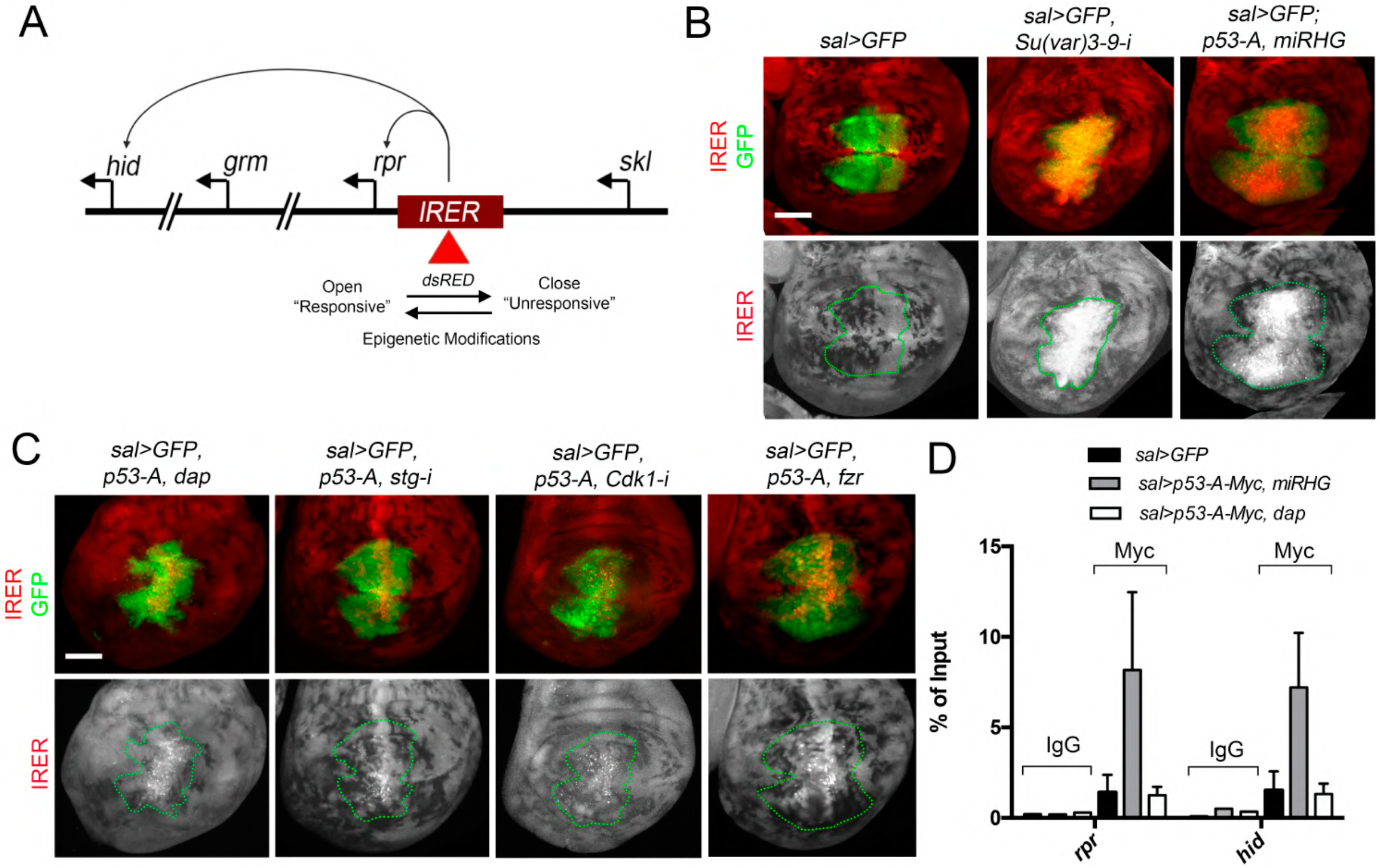
IRER activity and p53-A binding at the p53 ^RE^ of the pro-apoptotic genes *hid* and *rpr*. A) Schematic representation of the *Drosophila* H99 locus and the location of the irradiation responding enhancer region (IRER). The IRER contains a p53 ^RE^ and it is critical for apoptotic induction after irradiation. The accessibility of the IRER is subject to epigenetic regulation. An ubiquitin-DsRed reporter was inserted into IRER through homologous recombination ^69^. B) Third instar wing imaginal discs carrying the IRER {ubi-DsRed} cassette (red and white) and expressing the corresponding transgenes under the *sal>GFP* driver (green). Knockdown of Su(var)3-9 or the ectopic expression of *p53-A* along with the *miRHG* under the *sal-Gal4>GFP* driver, promote a strong response of the IRER{ubi-DsRed} cassette. Separate channel for the IRER is presented below each image with the *sal* domain marked with a green dotted line. C) Wing imaginal discs expressing *p53-A* and the different transgenes indicated in each panel under the *sal>GFP* driver. IRER activity is in red and GFP in green. Separate channel for the IRER is presented below each image with the *sal* domain marked with a green dotted line. D) Analysis of p53-A binding by chromatin immunoprecipitation experiments with anti-Myc or mock (IgG) at the p53 ^RE^ of the *hid* and *rpr* genes from wing imaginal discs of the following genotypes: *-sal>GFP* -*sal>GFP, p53-A (Myc), miRHG* -*sal>GFP, p53-A (Myc), dap*. Error bars represent SEM of three independent experimental replicates. In B and C the scale bar is 50 um. See also Figure S6.

To confirm that p53-A is unable to bind to the regulatory regions of the pro-apoptotic genes *rpr* and *hid* in cell cycle arrested cells, we performed a chromatin immunoprecipitation (ChIP) assay. We used negative control imaginal discs (*sal>GFP*) and discs that expressed *p53-A* tagged with Myc in proliferating (*sal>GFP, p53-A-Myc, miRHG*) and cell cycle arrested cells (*sal>GFP, p53-A-Myc, dap*) (Fig. 6D). We expressed the *miRHG* along with *p53-A* in the positive control proliferating discs to reduce the number of apoptotic cells that could interfere with the ChIP assay. Strong chromatin enrichment was observed at the p53^RE^ of the *rpr* and *hid* genes when *p53-A* was overexpressed in proliferating cells, however, this enrichment was not observed in cell cycle arrested cells (Fig. 6D).

Altogether, our results demonstrate that p53-A binding to the regulatory regions of the pro-apoptotic genes is compromised in cell cycle arrested cells.

### p53 apoptotic response is controlled by the G2/M promoting factor Cdk1

Our detailed analysis of the induction of cell death after IR or after *p53-A* overexpression in cell cycle blocked cells, suggests that p53-A apoptotic activity is regulated at the G2/M phase. Progression from G2 to M requires the activation of the Cdk1/CycB complex by the phosphatase Stg that removes the inhibitory phosphates ^70^. Importantly, we have shown that depletion of Stg, Cdk1 or *fzr* overexpression strongly abolished both IR and p53-A induced apoptosis. All these results suggest that Cdk1 active complexes may regulate p53-A pro-apoptotic function. To test this hypothesis, we decided to induce Cdk1 activity by expression of *stg* in the posterior compartment and test whether IR induced-apoptosis is increased. In control discs without IR, the temporal expression of *stg* for 24 hrs in the posterior compartment had almost no effect on apoptosis (Fig. 7A). Remarkably, in wing discs from the same genotype dissected 3 hrs after IR, the expression of *stg* did not rescue the mitotic arrest, however, significantly increased the number of apoptotic cells when compared to an irradiated control disc (Fig. 7B and C). As a complementary experiment, we forced G1/S transition by the overexpression of *CycE* in the posterior compartment to increase the proportion of cells in G2 and monitor the apoptotic response after IR ^45,51^. The restricted expression of *CycE* in non-irradiated discs for 24 hrs induced a considerable amount of cell death in the posterior compartment (Fig. 7A). Nevertheless, in irradiated discs the temporal expression of *CycE* increased the apoptotic response when compared to a wild type irradiated control disc (Fig. 7B and C). Importantly, the increase of apoptosis induced by *stg* or *CycE* overexpression in irradiated discs is dependent on p53 activity (Fig. 7D and E).

**Figure 7:**
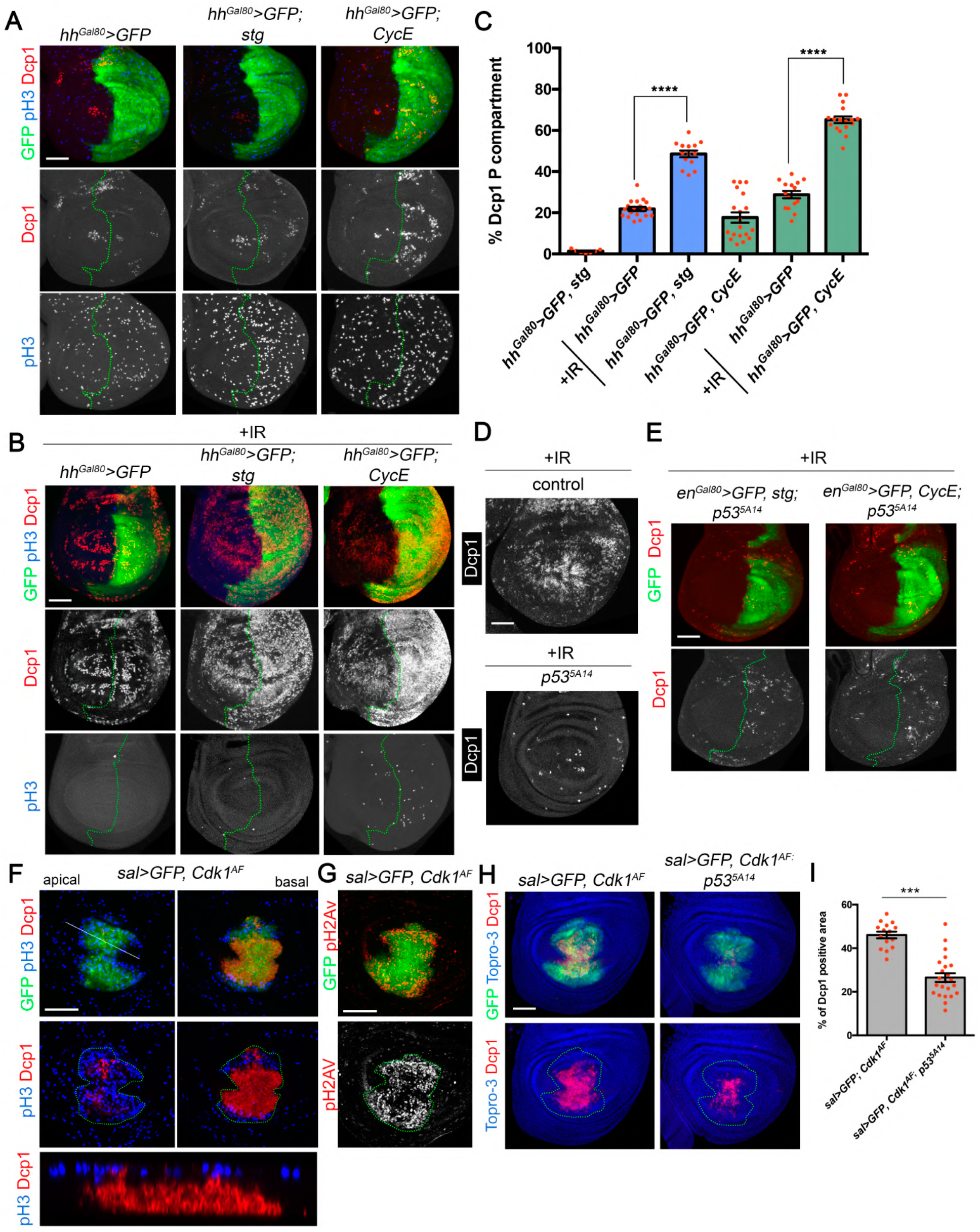
The G2/M promoting factor Cdk1 regulates p53 apoptotic response after IR. A) Third instar wing imaginal discs expressing the corresponding transgene for 24 hrs under the *hh-Gal4, tub-Gal80*^*ts*^ (*hh*^*Gal80*^*>*) and stained for Dcp1 (red), pH3 (blue) and GFP (green). Separate channels for Dcp1 and pH3 are shown. The antero-posterior compartment boundary is marked by a green dotted line. B) Third instar wing imaginal discs expressing the indicated transgenes for 24 hrs as in A, subjected to IR and dissected 3 hrs after treatment. Discs were stained for Dcp1 (red), pH3 (blue) and GFP (green). Separate channels for Dcp1 and pH3 are shown. The antero-posterior compartment boundary is marked by a green dotted line. C) Quantification of Dcp1 staining in the *hh* domain (posterior compartment) of wing imaginal discs from the genotypes and treatments described in A and B. Error bars indicate SEM. n>14 disc per genotype except for *hh*^*Gal80*^*>GFP, stg* where n=7. Statistically significant differences based on Student’s t test are indicated: ****P < 0,0001. D) Control and *p53*^*5A14*^ mutant wing discs from IR larvae and dissected 3 hrs later. Dcp1 staining is in white. E) Wing imaginal discs expressing the indicated transgenes under the *en-Gal4, tub-Gal80*^*ts*^ (*en*^*Gal80*^*>*) for 24 hrs in a *p53*^*5A14*^ mutant background. Third instar larvae were subjected to IR and dissected 3 hrs later. Wing discs were stained for Dcp1 (red and white) and GFP (green). Separate channels for Dcp1 are shown. The antero-posterior compartment boundary is marked by a green dotted line. F-G) Expression of the non-inhibitable version of Cdk1(*Cdk1*^*AF*^) in the *sal* domain of third instar wing discs stained for pH3 (Blue), Dcp1 (red) and GFP (green) in F and for pH2AV (red and white) in G. An apical, basal and Z-section are shown in F. The *sal* domain is marked by a green dotted line in F and G. Separate channel for pH2AV staining is shown in G. H) Third instar wing imaginal discs expressing the *Cdk1*^*AF*^ transgene under the *sal>GFP* driver in a control and a *p53*^*5A14*^ mutant background. Imaginal discs were stained for Dcp1 (red), Topro-3 (blue) and GFP (green). The *sal* domain is marked by a green dotted line. I) Dcp1 staining quantification in the *sal* domain of wing imaginal discs from the genotypes indicated and described in H. Error bars indicate SEM. n>15 disc per genotype. Statistically significant differences based on Student’s t test are indicated: ***P < 0,001. In A, B, D, E, F, G and H the scale bar is 50 um.

The expression of a non-inhibitable version of Cdk1 (T14A, Y15F, also called Cdk1^AF^) force cells to enter mitosis and to bypass the G2/M checkpoint ^71^. In addition, we observed that *Cdk1*^*AF*^ expression in the *sal* domain induced a strong apoptotic response, detected mostly in pH3 negative cells, that is associated with an increase of ATM/ATR activity visualized by pH2Av staining (Fig. 7F and G). This ectopic apoptotic induction is partially dependent of p53 activity, as Dcp1 staining was significantly reduced in the *p53*^*5A14*^ mutant background of *sal> Cdk1*^*AF*^ discs (Fig. 7H and I). These results support the role of Cdk1 as a regulator of p53 pro-apoptotic activity in G2/M.

To study the molecular connection between p53 and Cdk1 in more detail, we tested whether p53 and Cdk1 physically interact and its functional relevance in regulating the apoptotic response. We used the bimolecular fluorescence complementation assay (BiFC) to evaluate the direct physical interaction between p53 and Cdk1 in wing imaginal cells ^72,73^. This method is based on the reconstitution of the Venus fluorescent protein when two non-fluorescent Venus fragments fused to the proteins of interest (in this case VC-Cdk1 and VN-p53-A) are brought together in the cell (Fig. 8A). The expression of each individual construct in the wing imaginal cells by the *sal-Gal4* driver did not show any BiFC signal, however when VC-Cdk1 and VN-p53-A were co-expressed, strong nuclear Venus signal could be detected (Fig. 8C and D). Importantly, this signal is specific as the expression of two unrelated proteins failed to complement p53-A or Cdk1 as strong as the p53-A/Cdk1 complex (Fig. S7). In addition, the expression of a “cold” competitive partner of Cdk1 (Cdk1-HA) led to a decrease to background levels of the Cdk1/p53-A BiFC signal (Fig. S7).

**Figure 8:**
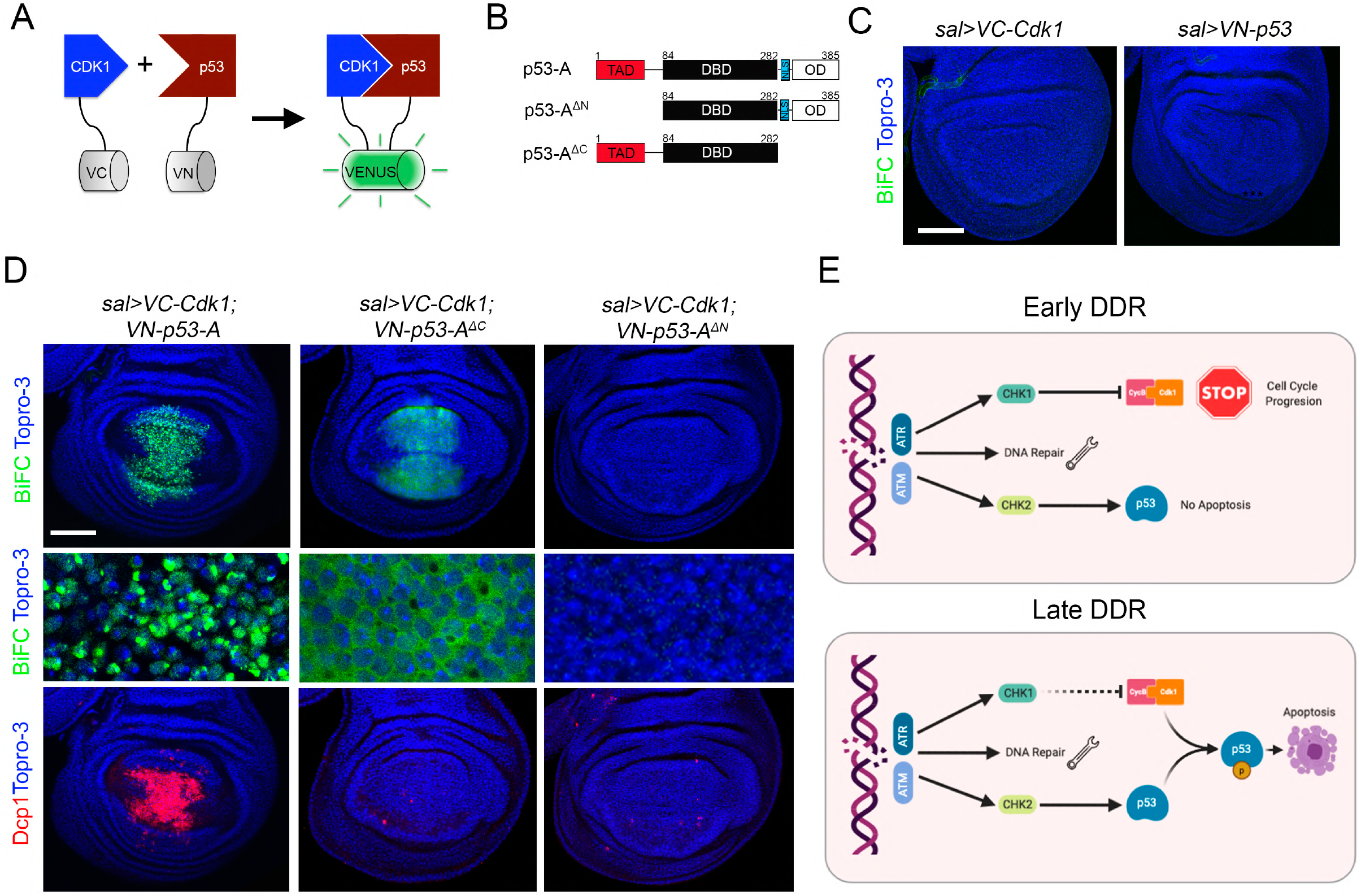
p53 interacts with Cdk1. A) Cartoon illustrating the BiFC principle. No fluorescent N-and C-terminal fragments of the GFP variant Venus are fused to p53 (VN-p53) and Cdk1 (VC-Cdk1). When these proteins are co-expressed inside cells juxtaposes the Venus fragments resulting in structural complementation and green fluorescence, enabling the direct visualization of protein interactions in living cells. B) Scheme of the different p53-A protein versions generated for the BiFC analysis. Amino acid positions defining p53 domains are indicated: transactivation domain (TAD, in red), DNA-binding domain (DBD, in black), nuclear localization signal (NLS, in blue) and oligomerization domain (OD, in white). C) Third instar wing imaginal discs expressing only *VC-Cdk1* or *VN-p53-A* under the *sal-Gal4* driver did not show any BiFC signal (green). Topro-3 staining marks nuclei in blue. D) BiFC analysis by the co-expression of *VN-p53-A, VN-p53-A*^*ΔC*^ and *VN-p53-A*^*ΔN*^ with *VC-Cdk1* under the *sal-Gal4* driver. Note that *p53-A-VN* and *VC-Cdk1* induced a strong nuclear BiFC signal (green) and the induction of apoptosis (Dcp1, red). The expression of the C-terminal deletion, *VN-p53-A*^*ΔC*^ with *VC-Cdk1* generates a cytoplasmic BiFC signal, but failed to induce cell death. In contrast, neither BiFC signal nor apoptosis was observed by the co-expression of *VC-Cdk1* with the N-terminal deletion of *p53-A* (*VN-p53-A*^*ΔN*^). A higher magnification of a region in the *sal* domain is presented. E) Simplified representation of the DNA damage response (DDR) model. DNA damage triggers the key signal transducers ATR and ATM kinases that activate the downstream effectors CHK1 and CHK2. CHK1 stop cell-cycle progression through the inactivation of Cdk1 and activates the DNA damage repair mechanisms. CHK2 induces the activation of p53, however, the pro-apoptotic activity of p53 is blocked in cell cycle arrested cells due to the presence of inactive Cdk1/CycB complexes. Some hours later, the G2/M blockage is progressively lifted, allowing the phosphorylation of p53 by active Cdk1/CycB and promoting p53 pro-apoptotic activity in cells unable to repair their DNA lesions. In C and D the scale bar is 50 um. See also Figure S7.

Next, we investigated which p53 domain is responsible for its interaction with Cdk1. We created p53 N- and C-terminal deletions for the transactivation (TAD) or the nuclear localization signal (NLS) and oligomerization domains (OD), respectively (Fig. 8B). While the C-terminal deletion (VN-p53-A^ΔC^) maintains its ability to interact with VC-Cdk1 as visualized by BiFC signal, this complex is mainly observed in the cytoplasm and it is unable to trigger apoptosis (Fig. 8D). In contrast, deletion of the N-terminal domain (VN-p53-A^ΔN^) strongly abolished the p53-A/Cdk1 BiFC signal and the apoptotic induction (Fig. 8D).

Together, these results indicate that p53 can interact directly with Cdk1 through the TAD.

## Discussion

### Cell cycle arrest blocks IR-induced apoptosis

In this work we study the connection between cell cycle progression and apoptotic induction after DNA damage. We demonstrate that IR-induced p53 pro-apoptotic activity is compromised in cell cycle stalled or endocycle-induced cells, as the p53 target gene *hid* or the effector caspase Dcp1 are not properly activated. In addition, the induction of the JNK pathway in response to IR is also blocked in these conditions. The JNK pathway is initially triggered in response to IR in a p53-dependent manner ^63^, but also in a late p53-independent way that contributes to the reduction of aneuploid cells generated following IR ^29,30,63^. Our results suggest that the activation of the JNK pathway is blocked in arrested cells due to defects on p53 activity and to the reduction in genome instability due to the blockage of mitosis ^29^.

Several mechanisms have been proposed to explain the differential apoptotic sensitivity of DNA damage cells depending on their proliferating status ^36,38,42^. This includes the epigenetic silencing of the regulatory regions of the pro-apoptotic genes and the proteasome-dependent degradation of p53 ^41,42^. Our results in wing disc cells demonstrate that p53 protein levels are comparable in cycling and experimentally arrested cells. In addition, using a sensor for the chromatin accessibility of the irradiation responsive enhancer region (IRER), we failed to observe a change in its activity in induced cell cycle arrested cells. However, we did find that p53 ability to induce an open chromatin structure of the IRER is strongly blocked in cell cycle arrested cells due to defects on p53 binding to the pro-apoptotic p53^RE^. Accordingly, p53 binding at the p53^RE^ of *rpr* and *hid* and the activation of the apoptotic program are also lost in a postmitotic tissue such as the adult heads when compared to a proliferating embryo ^37^. Our results are in accordance with the proposed role of p53 as a pioneer factor binding to p53^RE^ within inaccessible chromatin and inducing chromatin changes in response to DNA damage ^74-76^.

### Cdk1 connects the cell cycle with the apoptotic program

In human cells p53 is specifically phosphorylated by cdc2/Cdk1. This phosphorylation alters the binding site preference and enhances the sequence specificity binding of p53 to its target genes promoting their transcription *in vitro*^77-80^. Our results show that p53 is unable to bind to the p53^RE^ of the pro-apoptotic genes and to transcriptionally regulate their expression in cells with inactive Cdk1 complexes. Moreover, using BiFC we demonstrated that p53 physically interacts through its TAD domain with Cdk1. All these experiments suggest that after DNA damage, p53 requires an active Cdk1/CycB complex to mediate its pro-apoptotic activity. It would be interesting to study how Cdk1/CycB affects the p53 transcriptional output at a molecular level. One possibility is that p53 phosphorylation by active Cdk1 enables the selection of specific targets linking cell cycle progression to p53 transcriptional output. Supporting this hypothesis, it has been shown in *Drosophila* p53 regulates different DNA damage programs in a postmitotic tissue compared to a proliferating one through the differential p53 binding in these tissues ^37^.

### Life vs death decisions after DNA damage

Cells sensing DNA damage are faced with antagonizing responses such as cell cycle arrest and DNA repair or the induction of apoptosis. These “life or death” cell fate decisions are determined by a number of factors that include the activation of the DNA repair mechanisms, damage tolerance, the proliferation state of the cell or the cellular developmental context ^81^. Given the central role of p53 in the control of pro-survival and pro-apoptosis target genes, the precise regulation of p53 activity is essential for the appropriate response after genotoxic stress ^82-85^. Defects in the ability to trigger and coordinate any of these responses are a major cause of cancer development. Our results suggests a model where the connection between cell cycle progression, through the regulation of Cdk1/CycB activity, and the pro-apoptotic function of p53 allow cells with DNA damage to be protected from apoptotic induction when the DNA repair mechanisms are operational (Fig. 8E). After sensing DNA damage, cells activate the mitotic checkpoint, which in *Drosophila* arrest cells in G2/M, allowing time for the DNA repair mechanisms. This mitotic delay is dependent on ATR/Mei41 and Chk1/Grp that transiently downregulates Stg and therefore Cdk1 active levels ^20,24^. At the same time, in response to DNA damage ATM/Tefu phosphorylates Chk2/Mnk, which in turn activates p53 ^26^. Our results indicate that the pro-apoptotic function of p53 is blocked in cell cycle arrested cells due to the inactivation of the Cdk1/CycB complex. This cell cycle arrest gives cells an opportunity to repair their DNA lesions. Once the G2/M arrest is progressively lifted, cells unable to repair their DNA are sent to apoptosis through the activation of p53 by the ATM/Chk2 pathway and the presence of an active Cdk1/CycB complex.

Cell cycle arrest and apoptosis protection are hallmarks of cellular senescence ^86^. Our results suggest a possible common mechanism employed by cells to prevent apoptotic induction through the regulation of the cell cycle in different stress conditions such as DNA or tissue damage and senescence ^87,88^. Understanding the molecular basis of p53 apoptotic induction and its coordination with the cell cycle could have important implications in our understanding of tumor formation and cancer treatment.

## Acknowledgements

We thank Brian Calvi, Saeko Takada, Sonsoles Campuzano, Marco Milán, Hector Herranz, Ginés Morata, Samir Merabet, the Bloomington Stock Center, the Vienna Drosophila Resource Center and the Developmental Studies Hybridoma Bank for fly stocks and reagents. We specially thank the Confocal microscopy service at CBMSO, Eva Caminero and Mar Casado for fly injections, óscar Fernández-Capetillo for letting us use the Bioruptor and Ana Bermejo for qPCR assistance. We also thank Emilio Lecona and members of the lab for comments on the manuscript. This study was supported by grants from: FEDER/Ministerio de Ciencia e Innovación-Agencia Estatal de Investigación [No. PGC2018-095144-B-I00 to CE and BFU2014-54153-P to AB), the Fundación Ramón Areces and by institutional grant from Banco de Santander to the CBMSO.

## Author Contributions

Conceptualization, C.E. and A.B.; Methodology, M.R-L., R.G, A.P., A.B. and C.E.; Investigation, M.R-L., R.G, A.P., A.B. and C.E; Writing – Original Draft, C.E.; Writing –Review & Editing, C.E.; Funding Acquisition, C.E. and A.B.; Supervision, C.E and A.B.

## Declaration of Interests

The authors declare no competing interests.

## Materials and Methods

### Drosophila Strains

Reporters: Drice-based sensor (DBS-GFP) (Lopez-Baena), *hid-GFP, hid*^*20-10*^*-lacZ, JNK*^*REP*^*-RFP, IRER{ubi-DsRed}* and the Fly-FUCCI reporters *ubi-GFP-E2F11-230* and *ubi-mRFP1-NLS-CycB1-266 (ubi-flyFUCCI)* and *UAS-GFP-E2F11-230* and *UAS-mRFP1-NLS-CycB1-266* (Zeike et al, 2014). The Gal4 drivers: *ap-Gal4, tub-Gal80*^*ts*^ (*ap*^*Gal80*^*>*), *sal*^*EPv*^*-Gal4* (*sal>*), *dpp*^*blink*^*-Gal4* (*dpp>*), *hh-Gal4, UAS-GFP; tub-Gal80*^*ts*^ (*hh*^*Gal80*^*>*), *en-Gal4, UAS-GFP; tub-Gal80*^*ts*^ (*hh*^*Gal80*^*>*). The UAS lines: *UAS-dap, UAS-stg-i, UAS-Cdk1-i, UAS-fzr, UAS-CycE-i, UAS-E2f1-i, UAS-Rbf*^*280*^, *UAS-CycA-i, UAS-mre-11-i, UAS-okra-i, UAS-tefu-i, UAS-Dcr-2, UAS-rpr, UAS-hid, UAS-hep*^*CA*^, *UAS-GFP, UAS-mcherry, UAS-p53-A-myc, UAS-Su(var)3-9-i, UAS-miRNA-RHG, UAS-stg, UAS-CycE, UAS-Cdk1*^*AF*^, *UAS-p53-VN, UAS-Cdk1-HA, UAS-dac-VN* and *UAS-abdm-VC*. The following mutant lines were used: *lig4*^*169*^, *mei41*^*D5*^, *mnk*^*p6*^ and *p53*^*5A14*^. *UAS-Dcr2* was used in combination with different RNAi lines to enhance message RNAi lines to enhance message knockdown.

### Temperature shifts experiments

The temporal expression of the different UAS-lines was restricted when needed using the Gal4/Gal80 ^ts^ UAS system ^89^. Briefly, embryos were collected for two days, maintained at the restrictive temperature (17°C) and then shifted to the permissive temperature (29°C) for the appropriated time prior dissection.

### Imaginal discs staining, image acquisition and analysis

Third instar larvae were dissected in PBS and fixed with 4% paraformaldehyde, 0,1% Deoxicholate and 0,1% Tritón X-100 in PBS for 25 minutes at room temperature. They were blocked in PBS, 1% BSA, 0.3% Triton for 1 hour, incubated with the primary antibody over night at 4 °C, washed four times in washing buffer (PBS 0.3% Triton) and incubated with the appropriate fluorescent secondary antibodies for 1.5 hours at room temperature in the dark. They were then washed and mounted in Vectashield (Cat# H-1000 RRID:AB_2336790) for confocal analysis.

TUNEL analysis was performed using In Situ ‘Cell Death Detection Kit’ (TMR Red) (#12156792 910) and ‘Tunel Dilution Buffer’ (#11966006001) kits, both from Roche.

All confocal images were obtained using a Leica LSM510 and LSM710 vertical confocal microscope. Multiple focal planes were obtained for each imaginal disc. Image treatment and analysis was performed using Fiji (https://fji.sc) and Adobe Photoshop software.

For the quantification of Dcp1 and Hid-GFP staining, a Z-maximal intensity projection was generated for each image and a high-intensity threshold was adjusted for each image. Then, we calculated the percentage of staining covered in the region of interest. The mitotic index was calculated as the average value of the ratio between the number of cells in mitosis (pH3-positive cells) and the area defined by the domains of expression the *sal>GFP*. The number of pH2AV foci was calculated similarly.

For the Fly-FUCCI cell quantification, third instar larvae of the *ap-Gal4; UAS-GFP-E2F11-230, UAS-mRFP1-NLS-CycB1-266* genotype were subjected to IR and dissected 1, 3 and 6 hrs after treatment. Non–irradiated larvae were used as control. Red, green or yellow cells were manually quantified. For each experiment, at least 5 wing imaginal discs were used to count an average of 500 cells per disc. The same region of the imaginal disc for each disc was selected for the quantification.

The number of discs analyzed in each experiment is given in the figure legends.

Statistical analysis was performed using Graph Pad Prism software (https://www.graphpad.com). The specific statistical test and the n used in each analysis are noted in the corresponding figure.

### Fluorescence Activated Cell Sorting (FACS)

50 wing discs were dissected from third instar larvae expressing the corresponding transgene under the *sal>GFP* driver. Larvae were incubated for 40 minutes at 28°C in 300 µl of trypsin solution (trypsin-EDTA, Sigma T4299) containing 1 µl of Hoechst (Hoechst 33342, Molecular Probes) in agitation. Trypsin digestion was stopped by the addition of 200 µl of 1% fetal bovine serum (FBS, Sigma 9665) in PBS. After centrifugation at 1500 g at 4°C for 5 minutes, cells were suspended in 200 µl of 1% FBS and cells were sorted by GFP expression using FACSCVantage SE (BD Biosciences). The cell cycle profiles of GFP-positive and GFP-negative cells were determined by Hoescht flourescence using a FACSCalibur flow cytometer (Becton Dickinson). The cell cycle profile was analysed using FloJo 7.5 software and Dean-Jett-Fox model.

### Ionizing Radiation (IR) treatments

Third instar larvae of the indicated genotypes were irradiated in X-ray machine Phillips MG102 at the standard dose of 4000R and dissected at the indicated times depending on the experiment and stated in each figure.

### Comet assay for wing imaginal disc cells

DNA strand breaks and alkali-labile sites were assessed via the alkaline version of the Comet assay. 60 wing imaginal discs cells were enzymatically individualized by incubation 20 min in TrypLE TM Express Enzyme (Thermo Fisher Scientific, Waltham, Massachusetts, USA) and stored at −80°C in freezing buffer (85.5 g/L sucrose and 50 mL/L DMSO prepared in 11.8 g/L citrate buffer at pH 7.6) until use.

An estimated 104 cells were embedded in 0.75% low melting point agarose (LMPA) and deposited on pre-coated slides with 1% agarose. Immediately after agarose solidification (10 min on ice), samples were incubated for 1 hr at 4°C in a cold lysis buffer (2.5 M NaCl, 100 mM EDTA, 10 mM Tris, 1% Triton X-100, pH10). The slides were then rinsed in 0.4 M Tris, pH 7.4. Subsequently, DNA was allowed to unwind for 40 min in the electrophoresis buffer (300 mM NaOH, 1 mM EDTA, pH > 13) and electrophoresis was carried out for 30 min at 25 V and 300 mA (0.73 V/cm). Slides were neutralized in 0.4 M Tris pH 7.4 and stained with 50 µL of GelRed (Thermo Fisher Scientific). Samples were examined with a Leica DMI 3000B microscope (Germany), equipped with an EL6000 compact light source and a 480–550 nm wide band excitation filter and a 590-nm cut-off filter. Scoring was carried out using the OpenComet plug-in for the image-processing platform ImageJ. A total of 250 randomly selected cells were analyzed per condition. The tail moment (tail intensity x length summed over the whole extent of the tail) was used to measure DNA damage.

### Bimolecular fluorescence complementation assay

We used an pUASTattB that have the N-terminal (VN: 1-173) and C-terminal (VC: 155-238) moieties of Venus cloned in Xho1 and Xba1 restriction sites. The coding region of p53-A and Cdk1 were PCR amplified from the GH11591 clone (BDGP) and LD38718 clone (BDGP), respectively. Inserts were cloned in Xho1-Xba sites into the pUASTattB VN or VC version including a five amino acids linker region. N and C terminal deletions of p53-A were cloned in a similar manner. To ensure similar expression levels, all UAS constructs were inserted into the same attP site (86Fb), except UAS-VC-Cdk1 that was also inserted in 51D. The sequence of all primers used in this study:

UAS-VN-p53-A:

Forward:

5’-CAGT**CTCGAG**<u>GGCGGCTCAGGCGGC</u>ATGTATATATCACAGCCAATGTCGTG GC-3’

Reverse: 5’-CAGT**TCTAGA**TCATGGCAGCTCGTAGGCACG-3’ UAS-VN-p53-A^ΔN (1-83)^ Forward: 5’-CAGT**CTCGAG**<u>GGCGGCTCAGGCGGC</u>ATGGAGAATCACAACATCGGTGG-3’

Reverse:

5’-CAGT**TCTAGA**TCATGGCAGCTCGTAGGCACG-3’ UAS-VN-p53-A^ΔC (283-385)^ Forward: 5’-

CAGT**CTCGAG**<u>GGCGGCTCAGGCGGC</u>ATGTATATATCACAGCCAATGTCGTG GC-3’

Reverse:

5’-CAGT**TCTAGA**TCAGGACTTGCGCTTCTTGCTATTGAGCTGGCG-3’ UAS-VC-Cdk1:

Forward: 5’-

CAGT**CTCGAG**<u>GGCGGCTCAGGCGGC</u>ATGGAGGATTTTGAGAAAATTG-3’ Reverse: 5’-CAGT**TCTAGA**TTAATTTCGAACTAAGCCCGATTGAAAAC −3’

For the initial BiFC analysis, we used the UAS-p53-VN flies available at FlyORF (F004757).

Visualization and quantification of the BiFC signal was done using identical parameters for image acquisition between the different genotypes and analyzed using Fiji.

### Chromatin Immunoprecipitation (ChIP) and quantitative Real-Time PCR assay

The wing imaginal discs of 100 larvae were dissected for each condition, performing three replicates per ChIP. The genotypes used for each ChIP are described in the figure legends and the main text. Larvae were fixed in FA Fix Solution (1.8% Formaldehyde, 50mM HEPES pH8, 1mM EDTA pH8, 100mM NaCl) for 25 minutes at RT. Then, the tissue was incubated with Quench Buffer (1X PBS, 0.125M Glycine, 0.01% Triton X-100) for 6 minutes at RT. Larvae were washed with buffer A (10mM HEPES pH8, 10mM EDTA pH8, 0.5mM EGTA pH8, 0.25% Triton X-100) and buffer B (10mM HEPES pH8, 200mM NaCl, 1mM EDTA pH8, 0.5 mM EGTA pH8, 0.01% Triton X-100) consecutively, 20 minutes each at 4°C.

Wing imaginal discs were dissected in Buffer B on ice. Later, discs were centrifuged at max speed for 3 minutes at 4°C. Collected disc pellet that was resuspended in buffer C (10mM HEPES pH8, 1mM EDTA pH8, 0.5mM EGTA pH8) supplemented with 1mM PMSF and 1X protease inhibitors cocktail (Roche #11873580001). The discs were homogenized in this medium before proceeding to sonication. The tissue was sonicated 20 cycles (30’’ ON/30’’ OFF), at high power and at 4°C using diagenode bioruptor sonicator. We removed 10% from the samples for INPUT. Samples were precleared with protein G Affinity Gel (Sigma-Aldrich #E3403) for 1 hr on rotator at 4°C, and then the chromatin was transferred to a fresh tube. The anti-myc Affinity gel (Sigma-Aldrich #E6654) and the protein G Affinity Gel (Sigma-Aldrich #E3403) as negative control of each Chip were blocked in 1X RIPA (140mM NaCl, 20 mM HEPES pH8, 2 mM EDTA pH8, 2% Glycerol, 2% Triton x-100, 0.2% DOC) supplemented with 100ug/mL salmon sperm DNA and 100ug/mL BSA overnight at 4°C.

Next day, we pelleted beads at 6000 rpm for 2 minutes and mixed the chromatin with the blocked beads. The samples were incubated for 4h at 4°C. Finally, the beads were washed four times in RIPA 1X for 5 minutes at 4°C, pellet at 6000 rpm for 2 minutes at 4°C. Beads were washed again in TE for 5 minutes at 4°C and pellet at 6000 rpm for 2 minutes at 4°C

Chromatin was eluted from beads in TE with 1% SDS and 0.1M NaHCO3 at 50°C. To reverse the crosslinks, we incubated the eluted material at 65°C overnight, processing the INPUT of each sample in parallel. The next day, we added 50 ug/ml of RNAase and incubated the samples 30 minutes at 37°C. Then, 20 ug of Proteinase K were added and incubated 55°C for 3h. We added 4 uL NaCl 5M and 4uL Tris 1M to each sample before purifying them by phenol/chloroform method. Finally, we added 1,25 ul of glycogen (Roche #10901393001), 25 ul of NaAC 3M pH5.2 and 550ul of ethanol 100% at −20°C ON. Samples were spin for 20 minutes at maximum speed and the pellet was twice with ethanol 70% at −20C and centrifuged again for 10 minutes and let the pellet dry at RT. The samples were resuspended in 30 uL of TE buffer and amplified by qPCR using GoTag qPCR Master Mix (Promega #A6001), using as amplicons the p53^RE^ for the *rpr* and *hid* genes. Results were quantified using the delta Ct method and presented as percentage of input.

The sequence of the primers used in this study: *rpr*-p53-RE:

Forward: 5’-CTACGTTTCCCAGACCCAAGAC-3’

Reverse: 5’-GTCTCCATCCAATTCCCATCTC-3’ *hid*-p53-RE:

Forward: 5’-ACTTTTGTTCTTTTCGCTTTGGAC-3’

Reverse: 5’-GATGACGAAATTCAAGCACACTCT-3’

## Supplemental Information

**Fig. S1:**
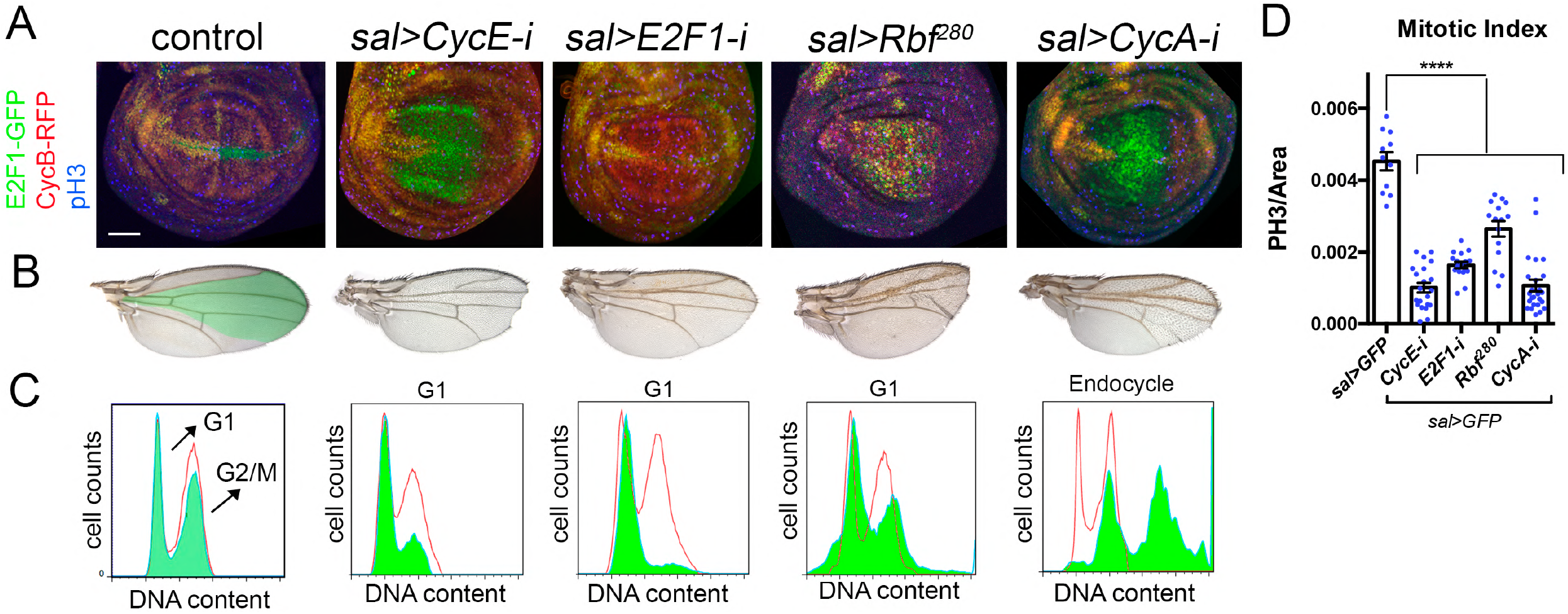
Analysis of cell cycle arrest tools. A) Cell cycle perturbations by the expression of *CycE-i, E2f1-i, Rbf*^*280*^ and *CycA-i* with the *sal-Gal4* (*sal>*) driver. The Fly-FUCCI system (*ubi-GFP-E2F11-230* and *ubi-mRFP1-NLS-CycB1-266*) and pH3 staining (blue) was used to visualize the cell cycle. B) Adult wing phenotypes of the experiments presented in A are shown. The *sal*domain is colored in green in the control. C) Cell cycle profiles of dissociated wing imaginal discs expressing the indicated cell-cycle regulators and GFP in the *sal* domain are shown. Red profiles correspond to control GFP negative cells and green profiles belong to GFP positive cells in control and cell cycle perturbed cells. D) Mitotic index measured as the number of pH3 positive cells per area in the *sal* domain of control (*sal>GFP*) and in cell cycle arrested cells and endocycle-induced cells. n>11 discs per genotype. Error bars indicate standard error of the man (SEM). **** P value <0,0001 by one-way ANOVA when compared the mean of each column with the mean of the control.

**Fig. S2:**
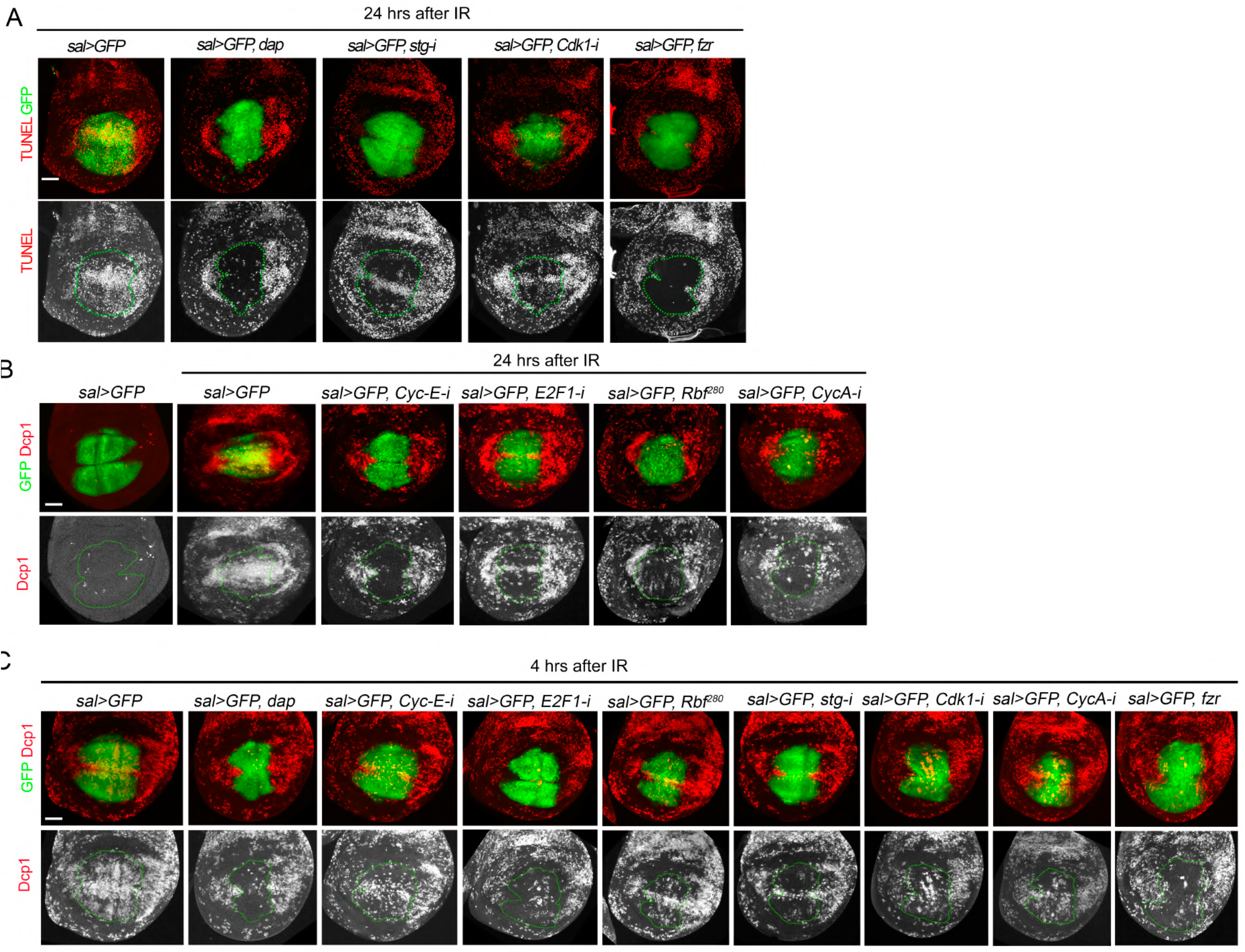
IR-induced apoptosis analysis in cell cycle arrested cells. A) GFP (green) and TUNEL staining (red and white) in wing imaginal discs expressing the indicated transgene by the *sal-Gal4* in irradiated discs and analyzed 24 hrs later. Below each panel, the TUNEL channel is shown and the *sal* domain is outlined by green dotted lines. B) Non-irradiated and irradiated wing imaginal discs expressing the indicated transgenes under the *sal-Gal4* driver. Discs were dissected 24 hrs after IR and stained for Dcp1 (red and white) and GFP (green). Below each panel, the Dcp1 channel is shown and the *sal* domain is outlined by green dotted lines. C) Wing imaginal discs expressing the indicated transgene by the *sal-Gal4* in control discs and irradiated discs analyzed 4 hrs later. GFP is in green and Dcp1 staining in red and white. The Dcp1 channel is shown and the *sal* domain is outlined by green dotted lines.

**Fig. S3:**
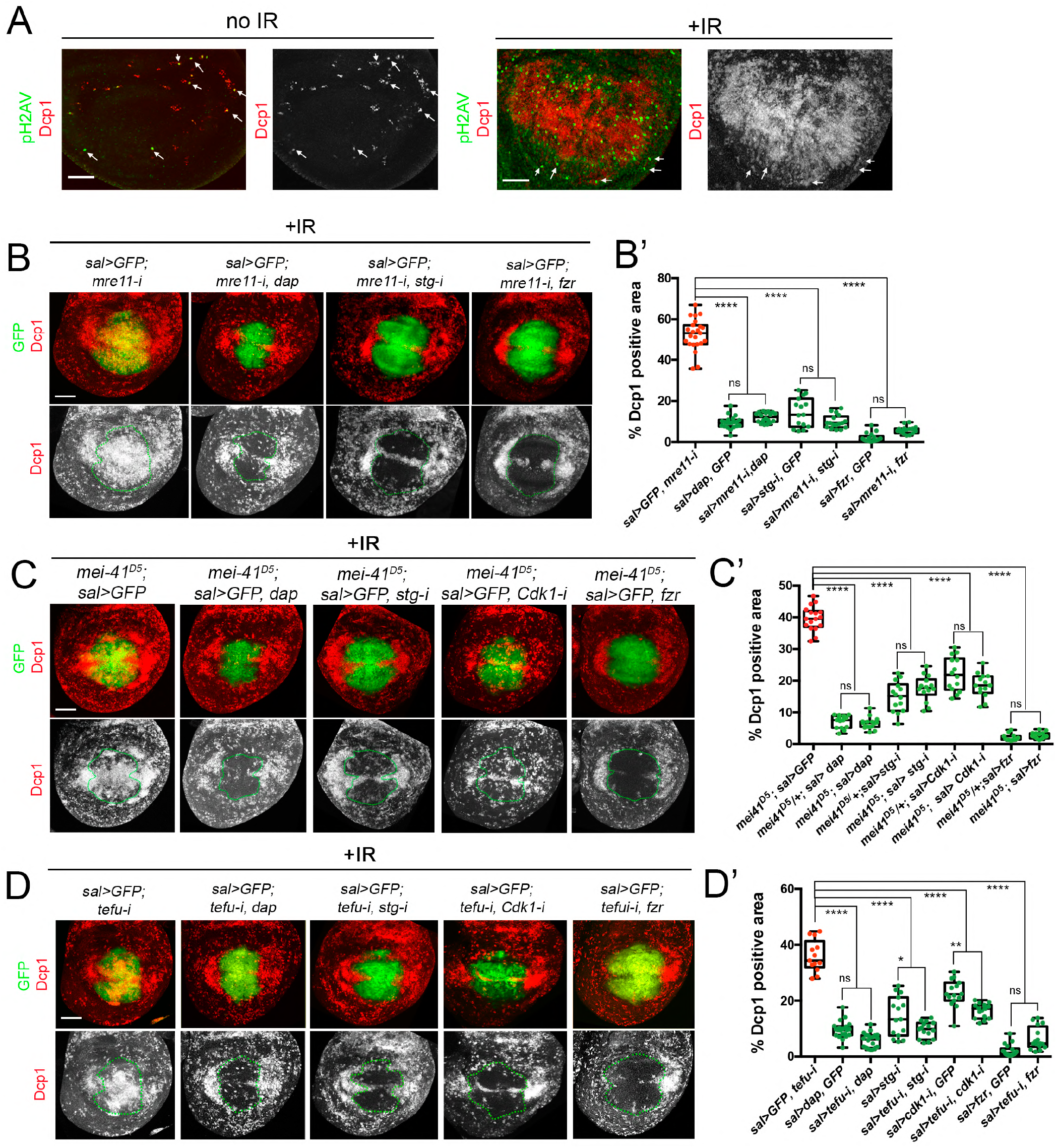
Apoptotic response after IR in cell cycle arrested and endocycle-induced cells that simultaneously knockdown DNA repair mechanisms. A) Strong pH2Av foci (green and arrows) are associated to cells with high levels of Dcp1 (red and white) in non-irradiated and irradiated wing imaginal discs. Separate channels for Dcp1 staining are shown. B) Irradiated wing imaginal discs analyzed 24 hrs later expressing the *mre11-i* line and the indicated transgenes by the *sal-Gal4, UAS-GFP* driver. *dap* arrest cells in G1, *stg-i* in G2 while *fzr* induces the endocycle.V GFP is green and Dcp1 staining is red or white. Below each panel, the Dcp1 channel is shown and the *sal* domain is outlined by green dotted lines. B’) Quantification of Dcp1 staining in the *sal* domain of wing imaginal discs of the corresponding genotypes presented. Error bars indicate the minimum and maximum point for each genotype. Individual wing discs measurements are shown. n>15 discs per genotype. **** P value <0,0001 by one-way ANOVA when compared the mean of each column with the mean of the corresponding control. ns, not significant. C) Irradiated wing imaginal discs analyzed 24 hrs later expressing the indicated transgenes by the *sal-Gal4, UAS-GFP* driver in a *mei-41*^*D5*^ mutant background. GFP is green and Dcp1 staining is red or white. Below each panel, the Dcp1 channel is shown and the *sal* domain is outlined by green dotted lines. C’) Quantification of Dcp1 staining in the *sal* domain of wing imaginal discs of the corresponding genotypes presented. Error bars indicate the minimum and maximum point for each genotype. Individual wing discs measurements are shown. n>15 discs per genotype. **** P value <0,0001 by one-way ANOVA when compared the mean of each column with the mean of the corresponding control. ns, not significant. D) Irradiated wing imaginal discs analyzed 24 hrs later expressing the *tefu-i* line and the indicated transgenes by the *sal-Gal4, UAS-GFP* driver. GFP is green and Dcp1 staining is red or white. Below each panel, the Dcp1 channel is shown and the *sal* domain is outlined by green dotted lines. D’) Quantification of Dcp1 staining in the *sal* domain of wing imaginal discs of the corresponding genotypes presented. Error bars indicate the minimum and maximum point for each genotype. Individual wing discs measurements are shown. n>15 discs per genotype. **** P value <0,0001, ** P value<0,01 and * P value<0,05 by one-way ANOVA when compared the mean of each column with the mean of the corresponding control. ns, not significant. Scale bar: 50 um.

**Fig. S4:**
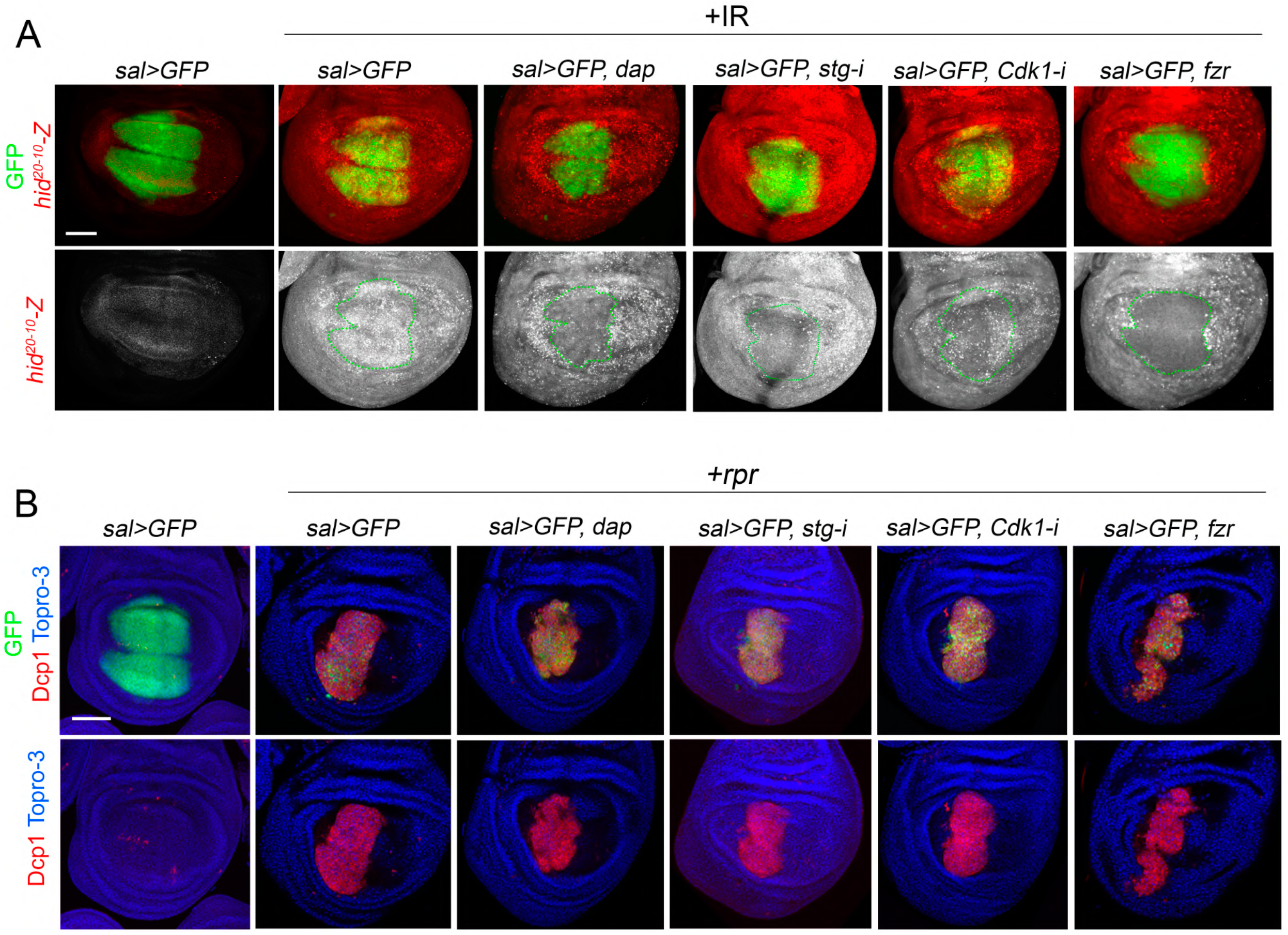
Analysis of Hid and Rpr activity in cell cycle arrested and endocycle-induced cells. A) Wing imaginal discs expressing the indicated transgenes under the *sal-gal4, UAS-GFP* driver and the *hid*^*20-10*^ regulatory region driving the *lacZ* reporter gene (*hid*^*20-10*^*-Z*) (red and white). Note that the *hid*^*20-10*^*-Z* is weakly active in the wing pouch in non-irradiated discs, however it is strongly activated 24 hrs after IR. *hid*^*20-10*^*-Z* separate channel is shown below each panel and the *sal* domain marked by green dotted lines. B) Dcp1 staining (red) in wing imaginal discs expressing *rpr* and the indicated transgenes under the *sal-Gal4, UAS-GFP* driver. GFP is in green, Dcp1 in red and Topro-3 in blue. Scale bar: 50 um.

**Fig. S5:**
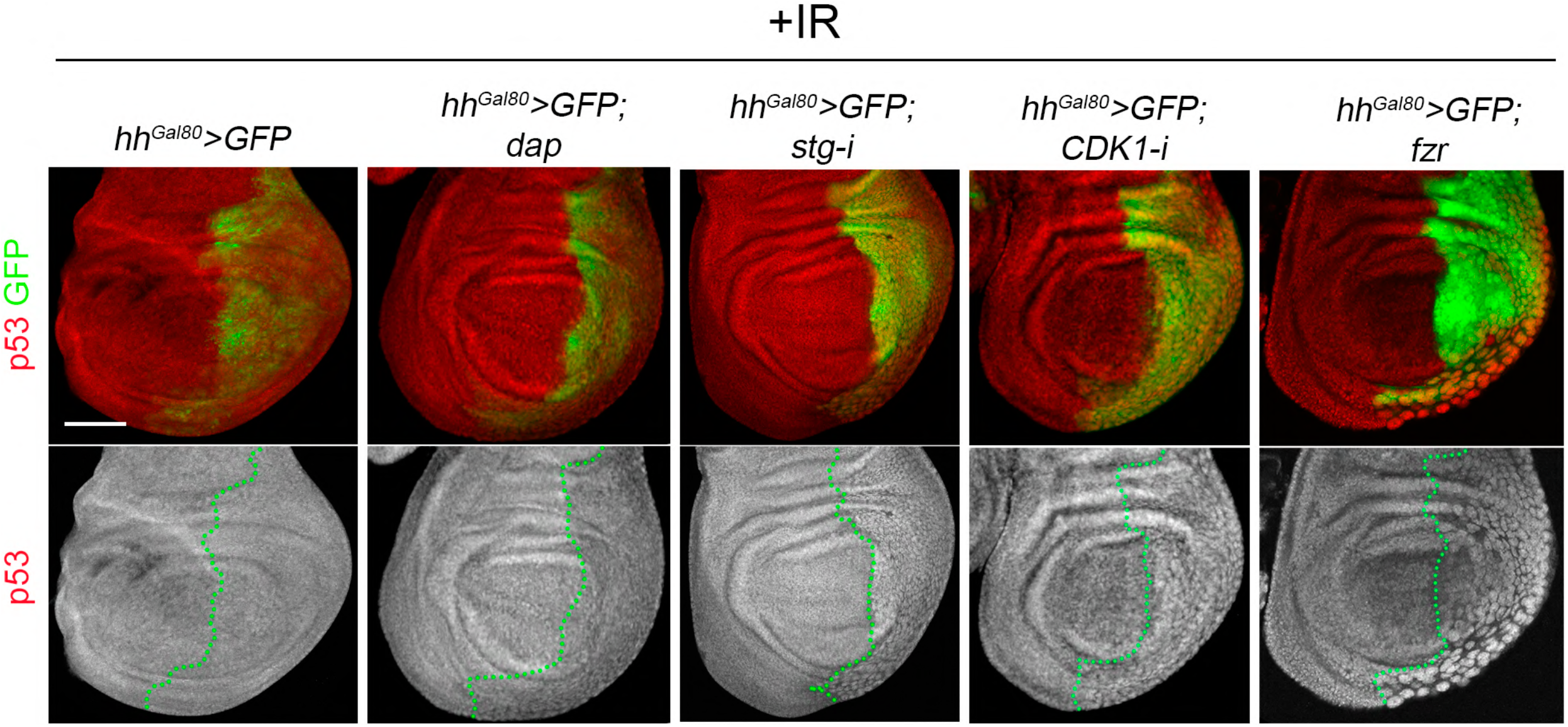
Analysis of p53 protein levels in cell cycle arrested and endocycle-induced cells of irradiated discs. Irradiated third instar wing imaginal discs expressing the indicated transgene with by the *hh-Gal4, tub-Gal80*^*ts*^ (*hh*^*Gal80*^*>*) driver and stained for p53 (red and white) and GFP (green). Separate channel for p53 is shown below each image. The antero-posterior compartment boundary is marked by a green dotted line. Larvae were kept at 17°C and shifted to 29°C for 24 hrs and wing discs were dissected 4 hrs after IR. Although a slight decrease of p53 protein levels could be observed in posterior cells expressing *dap*, no changes were detected for the other cell cycle modifications. Scale bar: 50 um.

**Fig. S6:**
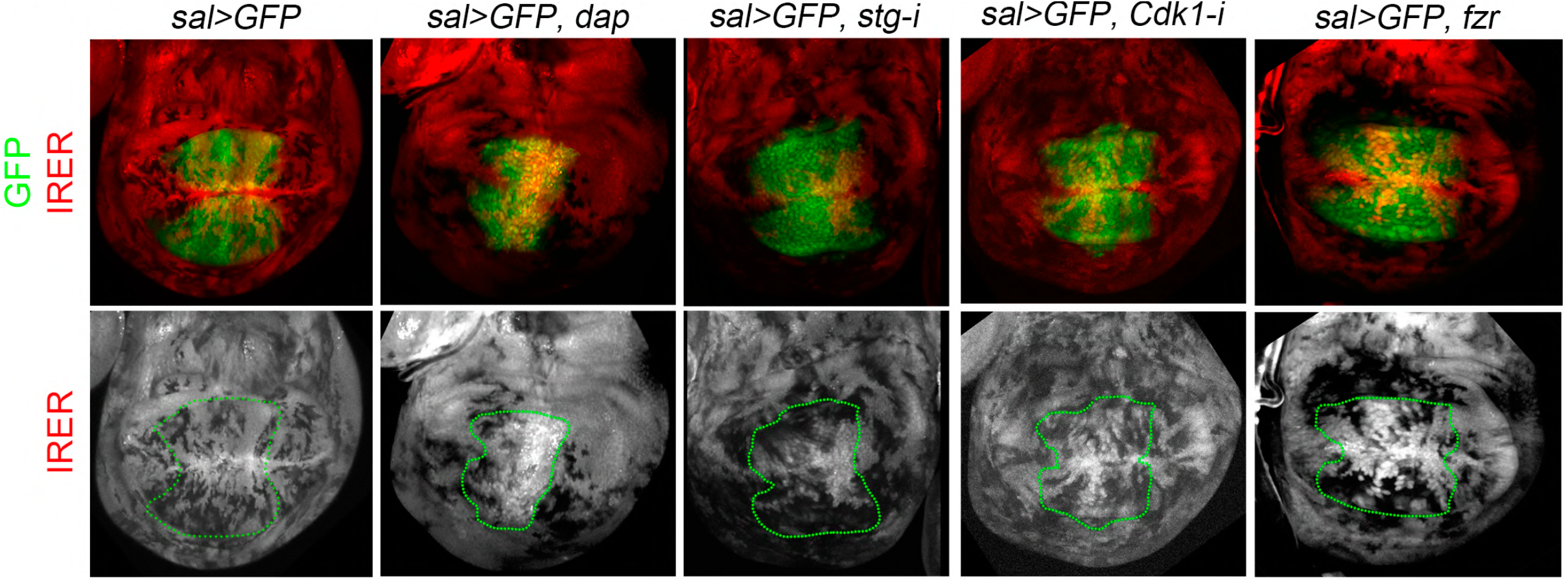
IRER activity in cell cycle arrested and endocycle-induced cells. Third instar wing imaginal discs carrying the IRER{ubi-DsRed} (red and white) cassette and expressing the corresponding transgenes under the *sal>GFP* driver (green). Separate channel for the IRER{ubi-DsRed} (white) is presented below each image with the *sal* domain marked with a green dotted line.

**Fig. S7:**
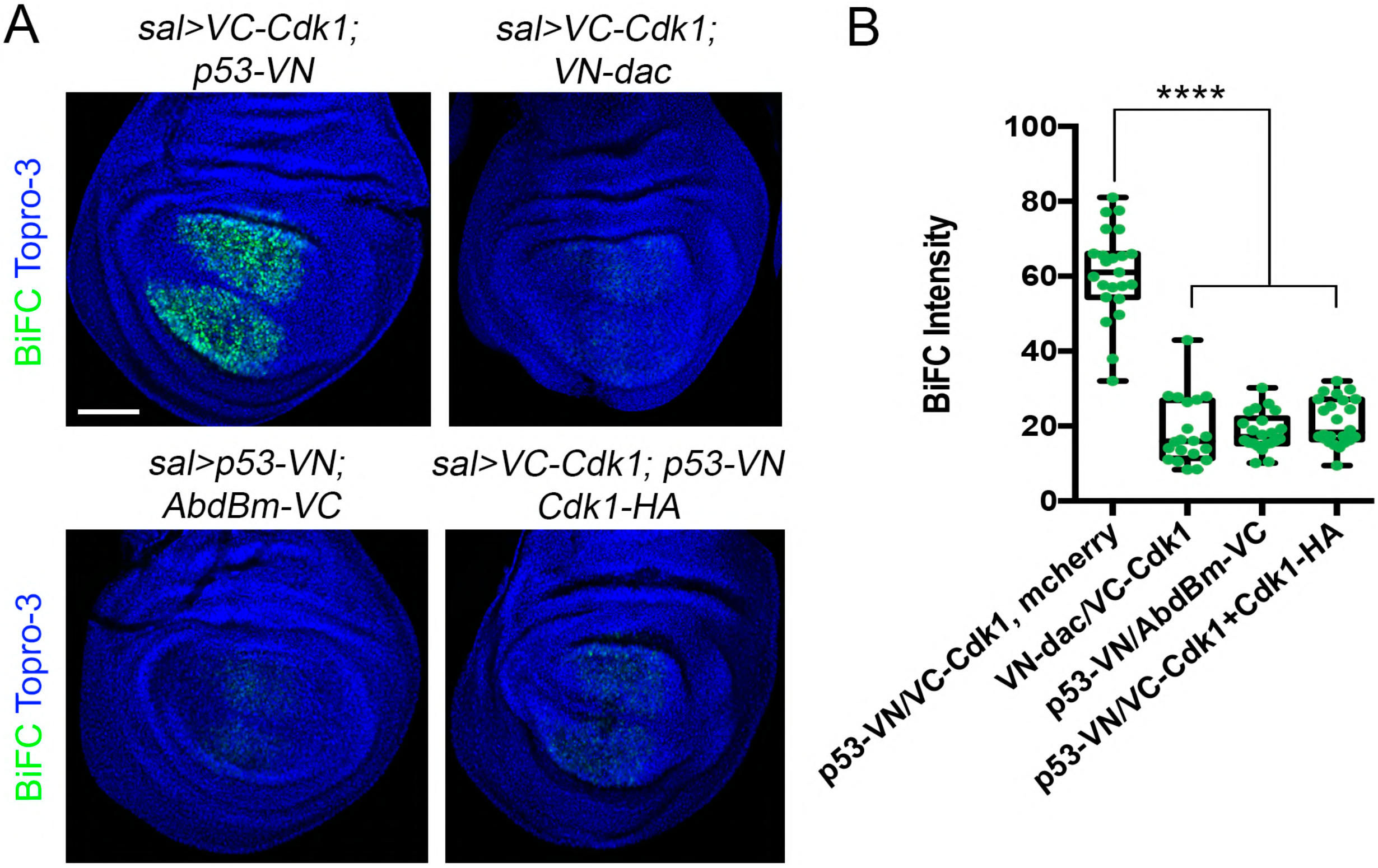
BiFC analysis of p53 interaction with Cdk1. A) Third instar wing imaginal discs expressing the indicated *VC* and *VN* fusion proteins under the *sal-Gal4* driver. Note that the strong BiCF signal is specific to the VC-Cdk1/p53-VN pair and that this signal is strongly reduced when a cold competitor (Cdk1-HA) is also co-expressed. B) Quantification of fluorescent signal resulting from the BiFC experiments. Data are represented as a boxplot showing each individual measurement. n>20 discs per genotype. Error bars indicate the minimum and maximum point for each genotype. **** P value <0,0001 by one-way ANOVA when compared the mean of each column with the mean of the corresponding control. All the images were taken keeping the same confocal settings.

## Notes

### Competing Interest Statement

The authors have declared no competing interest.

